# Functional comparisons of the virus sensor RIG-I from humans, the microbat *Myotis daubentonii*, and the megabat *Rousettus aegyptiacus*, and their response to SARS-CoV-2 infection

**DOI:** 10.1101/2023.02.08.527785

**Authors:** Andreas Schoen, Martin Hölzer, Marcel A. Müller, Christian Drosten, Manja Marz, Benjamin Lamp, Friedemann Weber

## Abstract

Bats (order *Chiroptera*) are a major reservoir for emerging and re-emerging zoonotic viruses. Their tolerance towards highly pathogenic human viruses led to the hypothesis that bats may possess an especially active antiviral interferon (IFN) system. Here, we cloned and functionally characterized the virus RNA sensor, Retinoic Acid-Inducible Gene-I (RIG-I), from the “microbat” *Myotis daubentonii* (suborder *Yangochiroptera*) and the “megabat” *Rousettus aegyptiacus* (suborder Yinpterochiroptera), and compared them to the human ortholog. Our data show that the overall sequence and domain organization is highly conserved and that all three RIG-I orthologs can mediate a similar IFN induction in response to viral RNA at 37° and 39°C, but not at 30°C. Like human RIG-I, bat RIG-Is were optimally activated by double stranded RNA containing a 5’-triphosphate end and required Mitochondrial Antiviral-Signalling Protein (MAVS) for antiviral signalling. Moreover, the RIG-I orthologs of humans and of *R. aegyptiacus*, but not of *M. daubentonii*, enable innate immune sensing of SARS-CoV-2 infection. Our results thus show that microbats and megabats express a RIG-I that is not substantially different from the human counterpart with respect to function, temperature dependency, antiviral signaling, and RNA ligand properties, and that human and megabat RIG-I are able to sense SARS-CoV-2 infection.

**IMPORTANCE:** A common hypothesis holds that bats (order *Chiroptera*) are outstanding reservoirs for zoonotic viruses because of a special antiviral interferon (IFN) system. However, functional studies about key components of the bat IFN system are rare. RIG-I is a cellular sensor for viral RNA signatures that activates the antiviral signalling chain to induce IFN. We cloned and functionally characterized RIG-I genes from representatives of the suborders *Yangochiroptera* and *Yinpterochiroptera*. The bat RIG-Is were conserved in their sequence and domain organization, and similar to human RIG-I in (i) mediating virus- and IFN-activated gene expression, (ii) antiviral signalling, (iii) temperature dependence, and (iv) recognition of RNA ligands. Moreover, RIG-I of *Rousettus aegyptiacus* (suborder *Yinpterochiroptera*) and of humans were found to recognize SARS-CoV-2 infection. Thus, members of both bat suborders encode RIG-Is that are comparable to their human counterpart. The ability of bats to harbour zoonotic viruses therefore seems due to other features.

## INTRODUCTION

Bats (order *Chiroptera*) are assumed to harbour more virus species than any other mammals including rodents (1-3). Several of the bat-borne pathogens (mostly RNA viruses) can cause severe disease in humans e.g. SARS coronaviruses 1 and 2 (3-5), or Ebola and Marburg viruses (6, 7). The taxonomic order *Chiroptera* was recently divided into the two suborders *Yangochiroptera* and *Yinpterochiroptera* (8), which largely, but not entirely, overlap with the previous division into microbats (having the ability of echolocation) and megabats (large fruit eating bats), respectively. Although members of both suborders can host highly pathogenic viruses, they rarely show clinical signs of disease, indicating an ability to tolerate and resist infection to an unprecedented level (9). Infection tolerance is proposed to be mediated by a dampening of pro-inflammatory responses that would otherwise lead to tissue damage (10-14). Infection resistance is supposed to be due to special features of the antiviral type I interferon (IFN) system, which is in bats under strong positive selection (15-19). In line with this, some *Yinpterochiroptera* “megabats” exhibit elevated base levels of type I IFNs, an expanded tissue distribution of the master IFN regulator IRF7, or diversified IFN gene loci (20-22), whereas *Myotis* “microbats” (*Yangochiroptera*) express uniquely high paralog numbers of the broadly antiviral IFN effectors Tetherin (BST2) and PKR (23-25).

Type I IFNs (IFN-*α*/*β*) are cytokines that constitute the first line of defence against viral infection (26). Several intra- and extracellular pattern recognition receptors (PRRs) are able to sense viral hallmark structures called pathogen-associated molecular patterns (PAMPs) like e.g. genomic RNA (27), and initiate a signal transduction chain that leads to the upregulation of IFN genes, first of all IFN-*β*. The RNA sensor Retinoic Acid Inducible Gene I (RIG-I) is one of the most important PRRs for virus infection (28). It has three major domains: an N-terminal domain encompassing two caspase recruitment domains (CARDs), a central DExD/H box RNA helicase domain, and a C-terminal domain involved in ligand binding and regulation (29, 30). RIG-I recognizes double-stranded RNA (dsRNA) structures - especially when containing a 5’triphosphate (5’ppp) moiety - as it is present in many RNA virus genomes (31-34). RNA ligand-bound RIG-I then undergoes a conformational change to expose the N-terminal CARD domains, thus enabling interaction with the signal adaptor Mitochondrial Antiviral Signalling Protein (MAVS) and the eventual activation of the IFN transcription factor, Interferon Regulatory Factor 3 (IRF3) (28). Newly produced and secreted IFN then binds to its cognate receptor on the cell surface to activate the expression of antivirally active interferon-stimulated genes (ISGs), thus establishing a virus-resistant cellular state.

So far, knowledge on RIG-I genes of bats is mostly limited to nucleotide sequences, sequence comparisons, genomic organization, and expression (35-40). The known bat RIG-I amino acid sequences (which are all bar one from *Yinpterochiroptera* “megabats”) exhibited approximately 70 to 90% similarity to RIG-I of other mammals, and suggested the conserved domain organization (37, 39). Basic mRNA levels of bat RIG-I were elevated in immune associated tissues eg. spleen, and could be stimulated by virus infection or the dsRNA mimetic poly I:C (35-37, 39-41), or type IFN (24, 42).

However, despite the importance of bats as major hosts for zoonotic viruses and the central role of RIG in antiviral defence, functional data on bat RIG-I proteins are entirely lacking. To fill this gap, we cloned and expressed RIG-I sequences from two bat species representing “microbats” and “megabats”, namely *Myotis daubentonii* (belonging to the *Yangochiroptera*) and *Rousettus aegyptiacus* (belonging to the *Yinpterochiroptera*), investigated their function in virus recognition, PAMP binding and IFN induction, and compared them to human RIG-I. Moreover, we tested the involvement of the bat RIG-Is in recognition of SARS-CoV-2 infection.

## RESULTS

### Bat cells induce an innate immune response upon virus infection

We have previously shown that cell lines derived from *M. daubentonii* (MyDaNi) and *R. aegyptiacus* (Ro6E-J) are capable of responding to exogenously added pan-species IFN (24, 43). Here, we investigated whether these cells could also produce endogenous antiviral IFN in response to virus infection. We employed a pair of genetically matched viruses that are either a strong IFN inducer (La Crosse virus lacking the IFN suppressor NSs; LACVΔNSs), or blocking IFN induction (La Crosse virus expressing the IFN suppressor NSs; wt LACV) (44). Replication of the NSs-deleted virus mutant LACVΔNSs is severely reduced in IFN competent cells and animals (45). Besides the two bat cell lines, we employed simian Vero E6 cells, which lack type I IFN expression (46) as a negative control, and the IFN competent human A549 cells as positive control. Each of the cell lines was infected with either of the two viruses at a low MOI (0,01) to ensure multistep growth, and incubated for 72 h. Fig. 1A shows that supernatants harvested from Vero E6 cells contained similar amounts of infectious viruses, indicating comparable replication. In the cell lines A549, MyDauNi and Ro6E-J, by contrast, yields of the LACV ΔNSs were reduced by approximately 2 orders of magnitude compared to the wt LACV. Thus, just as in the human positive control line A549, the two bat cell lines are restricting replication of the IFN-sensitive virus mutant, strongly suggesting that the bat cell lines have mounted an antiviral IFN response.

**Figure 1.**
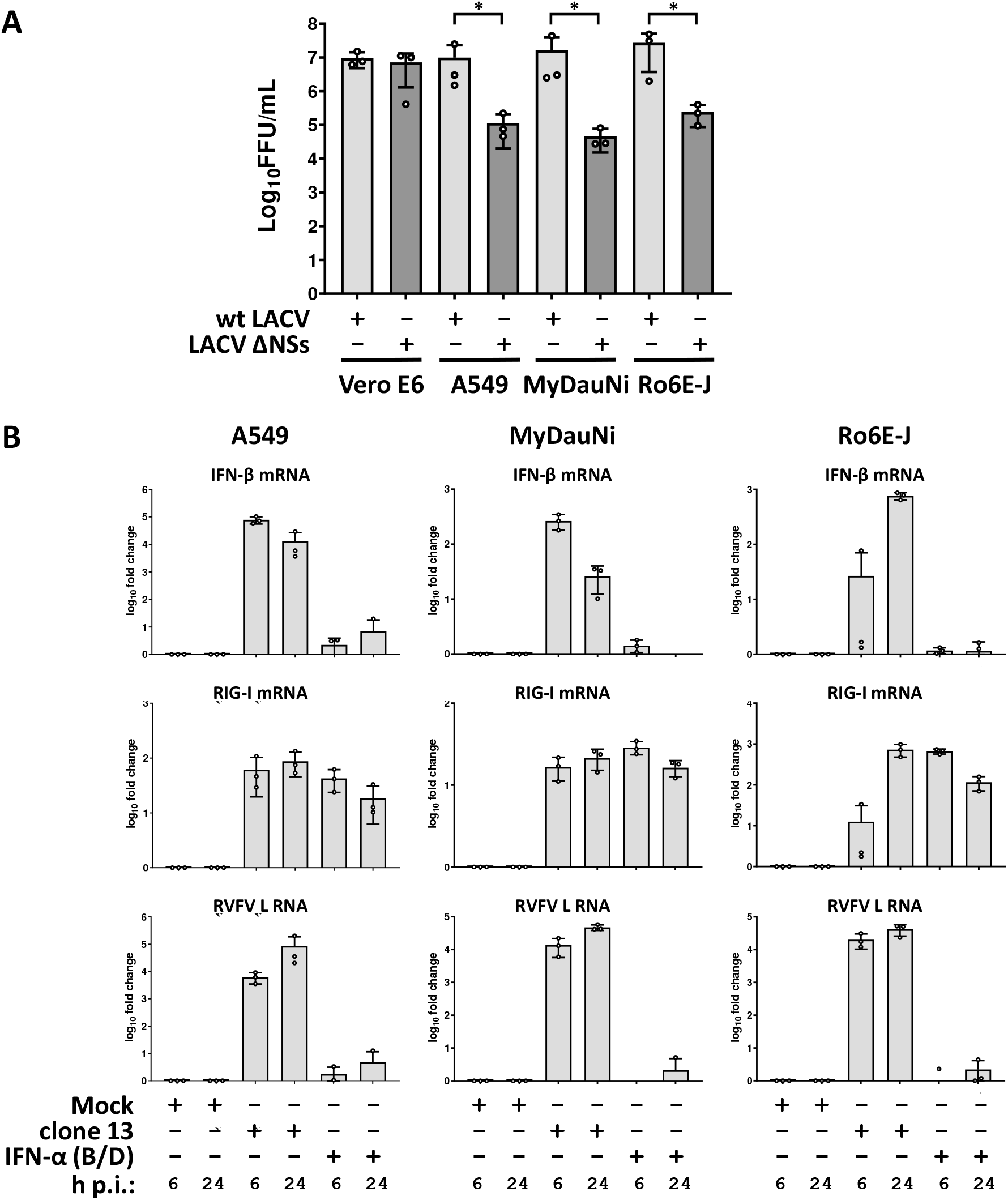
Type I Interferon competence of micro- and megabat cells lines. **A)** African green monkey kidney cells (Vero E6), human A549, *M. daubentonii* (MyDauNi) and *R. aegyptiacus* (Ro6E-J) cells were infected with wt LACV or the IFN-sensitive LACVΔNSs (MOI 0,01). After 72 h the supernatants were collected and the viral titers determined. The graphs show log10 titers, mean values and standard deviations from three independent replicates. **B)** A549, MyDauNi and Ro6E-J cells were either mock, pan-species IFN-α (B/D) (1000 U/ml) treated or infected with RVFV clone 13 (MOI 5) for 6 or 24 h, respectively. Expression of IFN-β, RIG-I, and RVFV L RNAs were monitored by RT-qPCR. The graphs show data points for log10 induction over mock, with mean values and standard deviations from three independent replicates. * p<0,05

To directly compare how the bat cells activate genes for endogenous IFN and ISGs (including RIG-I), we performed RT-qPCR analyses for RIG-I (DDX58) as well as for a series of other antiviral marker genes. IFN-β thereby represents exclusively virus-dependent genes (24, 47), similar to the chemokine CXCL10 (48) which is however also inducible by IFN-*γ* (49). Mx1 (MxA in humans) is established as an ISG that only reacts to IFN, but not directly to infection (50), whereas OAS1 can be induced by both virus infection and IFN (47). The A549, MyDaNi and Ro6E-J cells were either infected with the strong IFN-inducing Rift Valley fever virus (RVFV) mutant clone 13 (48), or treated with pan-species IFN-α. Total RNA samples were taken 6 and 24 h later, and tested for mRNA levels of the mentioned marker genes using RT-qPCR. Upon virus infection, all three cell lines induced IFN-β as expected, but with different kinetics (Fig. 1B). A549 and MyDauNi cells showed an initial peak at 6 h p.i. which then decreased to the later 24 h p.i. time point while Ro6E-J cells had a delayed IFN-β induction which peaked 24 h p.i. (Fig. 1B). Levels of viral RNA were however similar between the two bat cell lines, indicating true differences in IFN induction kinetics. RIG-I was upregulated in all three cell lines by clone 13 infection and by IFN-α treatment, but again the Ro6E-J cells reacted not as quickly to infection as the other two cell lines (see Fig. 1B). A similar Ro6E-J-specific pattern was observed for CXCL10, whereas all cells upregulated Mx1 and OAS1 in a similar manner (Fig. S1). These results indicate that the applied micro- and megabat cells are fully competent in launching an antiviral IFN response to viral infection or type I IFN treatment.

### Amino acid sequence comparison of bat RIG-Is

As a first step towards functional characterization of bat RIG-I orthologs, we assembled the full-length RIG-I sequence from *M. daubentonii*, using the data from our transcriptome studies (24, 43). Figure 2 shows the amino acid sequences alignment of the human, *M. daubentonii* and *R. aegyptiacus* RIG-I. Overall the sequence is well conserved, with 93,2 to 94,6 % sequence similarity between the different species. However, the bat orthologs exhibit two apparent differences to human RIG-I, namely a two-amino acid deletion at position 236-237 and an insertion of five amino acids at position 491-492. Whereas the short deletion is located in the hinge region between the CARDs and the helicase domain, the larger insertion locates in the Hel-2i helicase subdomain which is responsible for auto-inhibition of RIG-I in the absence of an RNA ligand (29). Moreover, *M. daubentonii* RIG-I has a one-amino acid deletion at position 195, which is also in the hinge region. Using the SMART database (http://smart.embl-heidelberg.de/), we identified the two N-terminal CARDs followed by a DEXD-like helicase domain, a helicase domain and the C-terminal regulatory domain as being conserved between all three RIG-Is (Sup. Fig. 3). Thus, while the overall sequence and domain organization is highly conserved, the two bat RIG-Is contain several indels.

**Figure 2.**
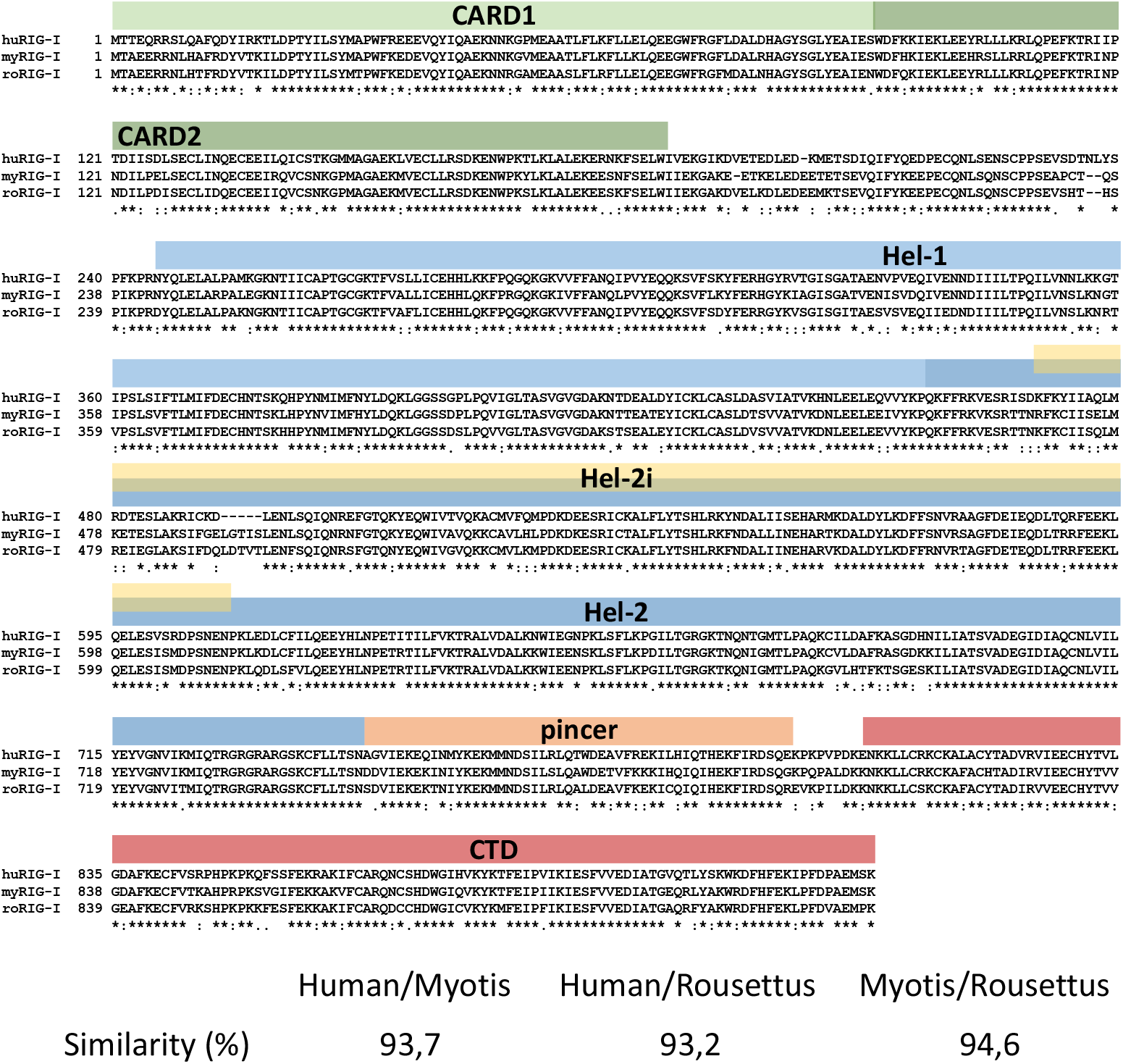
Amino acid sequence alignment for human, micro- and megabat RIG-I. Identical amino acid residues are marked with an asterisk (*), replacements with conserved (:) and semi-conserved residues with dots (.), and gaps with a dash (-). Above the alignment is the different domains of human RIG-I depicted, based on Kowalinski *et al*. (29). RIG-I sequences derived from human, *M. daubentonii*, and *R. aegyptiacus* are designated as huRIG-I, myRIG-I and roRIG-I, respectively.

**Figure 3.**
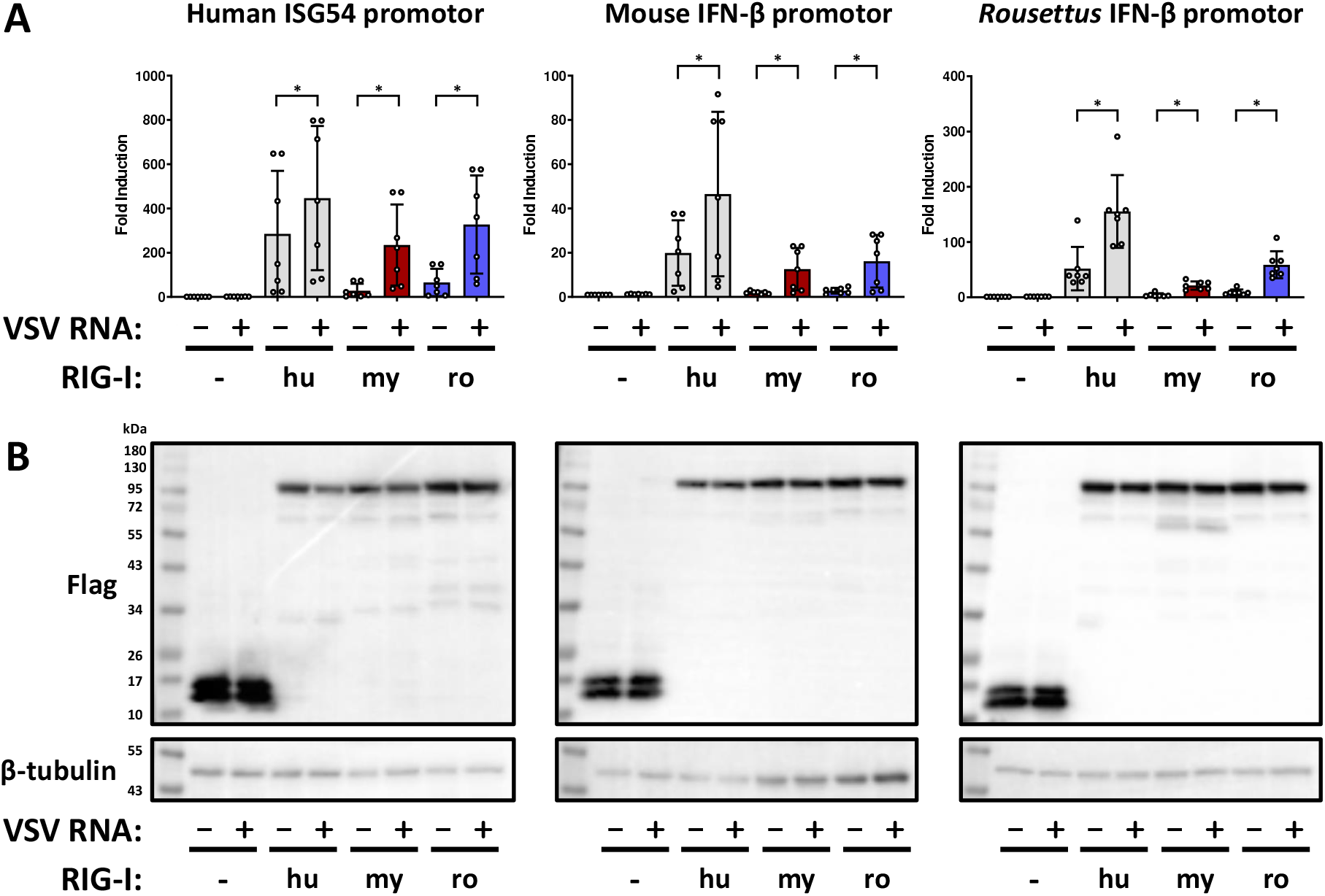
Transcomplementation of RIG-I-deficient human cells by bat RIG-I orthologs. **A)** HEK293 ΔRIG-I cells were transfected with plasmids encoding 3xFlag-ΔMx control protein (-), or 3xFlag-tagged hu-, my- or roRIG-I together with firefly-luciferase expressing plasmids under the control of the indicated innate immune promotors and *Renilla*-luciferase under SV40-control promotor. After 24 h, the cells were stimulated with VSV genomic RNA for 16 h. Upon harvesting the cells, the firefly/*Renilla* luciferase activities were measured. The graphs show data points for fold induction over the untreated negative control (column 1), with mean values and standard deviations from five independent replicates. **B)** Immunoblot analysis was performed with antibodies against the indicated antigens. Representative data from five independent experiments are shown. * p<0,05.

### Bat RIG-Is are functional

Attempts to knock down endogenous RIG-I in the bat cells or to find antibodies with sufficient cross-reactivity were unsuccessful (data not shown). We therefore constructed cDNA plasmids to express the three RIG-I orthologs, each equipped with an N-terminal 3×Flag epitope, in RIG-I-deficient cells. Then, we used reporter assays to test whether human HEK293 ΔRIG-I cells (33) could be transcomplemented with the cloned bat orthologs. The cells were transfected with the different RIG-I constructs (or a 3xFlag-control (CTRL) protein) together with plasmids encoding firefly luciferase under the control of the inducible human ISG54-promoter and a transfection control plasmids encoding *Renilla* luciferase under the control of the SV40-promoter. Since the ISG54 promoter is inducible by both virus infection and IFN, we also employed the exclusively virus-responsive IFN-β-promotors of mouse and of *R. aegyptiacus* in parallel. At 24 h after plasmid transfections, the inducible promoters were stimulated by supertransfection of the cells with the RIG-activating genome RNA of vesicular stomatitis virus (VSV). Following a further 16 h of incubation, firefly (inducible promoter) and *Renilla* (transfection baseline for normalization) luciferase activities were measured. The normalized data in figure 3A show an absence of inducible firefly luciferase activity in the CTRL-transfected samples, as expected, whereas expression of the human RIG-I alone already resulted in a 30-400 fold induction, depending on the promoter. As the CTRL-transfected cells did also not respond after VSV RNA transfection, any specific activity detected when RIG-Is are overexpressed is due to transcomplementation. When the cells expressing human RIG-I had been stimulated with VSV RNA, overall induction levels were 1,5-2,5 fold higher than in the unstimulated counterpart (see Fig. 3A). Unlike human RIG-I, the bat RIG-I orthologs did not exhibit background promoter induction, but in presence of VSV RNA they stimulated the promotors by 20-300 fold for *Myotis* and 20-400 fold for *Rousettus* (see Fig. 3A).

To make sure that the differences in induction levels were not due to differences in RIG-I expression, we analyzed the cell lysates by immunoblotting. All three RIG-Is from human, *Myotis* and *Rousettus* showed comparable levels and the expected apparent molecular weight of approximately 100 kDa (Fig. 3B). Moreover, we could also successfully transcomplement RIG-I^-/-^ mouse embryo fibroblasts with all three RIG-I orthologs, indicating that the RIG-I of humans and bats can rescue RIG-I deficiency irrespective of the species background (Fig. S4).

These results demonstrate that the RIG-I orthologs cloned from micro- and megabat cells can be stimulated by viral RNA and are able to initiate an antiviral signalling that results in the transactivation of virus-responsive promoters.

### Bat and human RIG-I show a similar induction pattern under different temperatures

The body temperature of bats is remarkably variable, from down to 11°C during sleep or hibernation to up to 41°C during flight (51). As the IFN system is known to be influenced by temperature (52), we wondered whether temperature might affect the activity of the bat RIG-I orthologs. To test this, we transcomplemented the HEK293 ΔRIG-I cells and allowed RIG-I expression for 24 h at 37 °C, but then placed the cells in incubators set to 30°C, 37°C or 39°C at 1 h before stimulation with VSV RNA, and kept them for another 16 h at the different temperatures. As shown in figure 4A, at 30°C none of the RIG-I orthologs could be specifically stimulated by VSV RNA. At 37°C, there was a robust activation as shown above, which was maintained at the temperature of 39°C. RIG-I expression levels were comparable at all temperatures, with a smaller unspecific band that is occasionally observed for *Myotis* RIG-I (Fig. 5B). These results indicated that the human and bat RIG-Is are functional at normal body temperature or higher, but not at 30°C.

**Figure 4.**
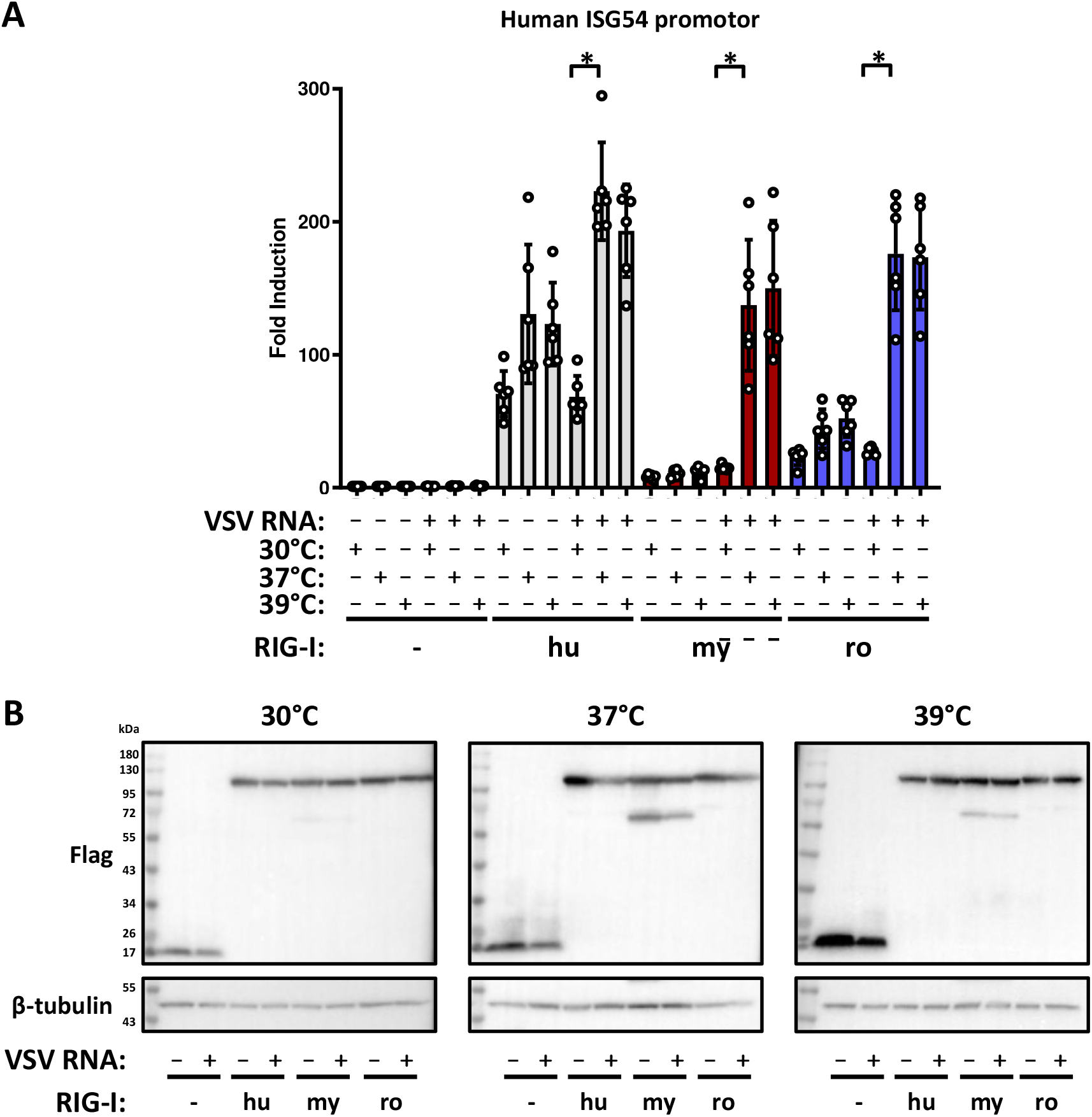
Temperature dependent IFN induction by exogenously expressed human and bat RIG-I orthologs. **A)** HEK293 ΔRIG-I cells were transfected, stimulated and assayed as described for figure 3, but incubated at different temperatures as indicated. Each graph shows the results from six independent replicates. **B)** Immunoblot analysis was performed with antibodies against the indicated antigens. Representative data from three independent experiments are shown. * p<0,05.

**Figure 5.**
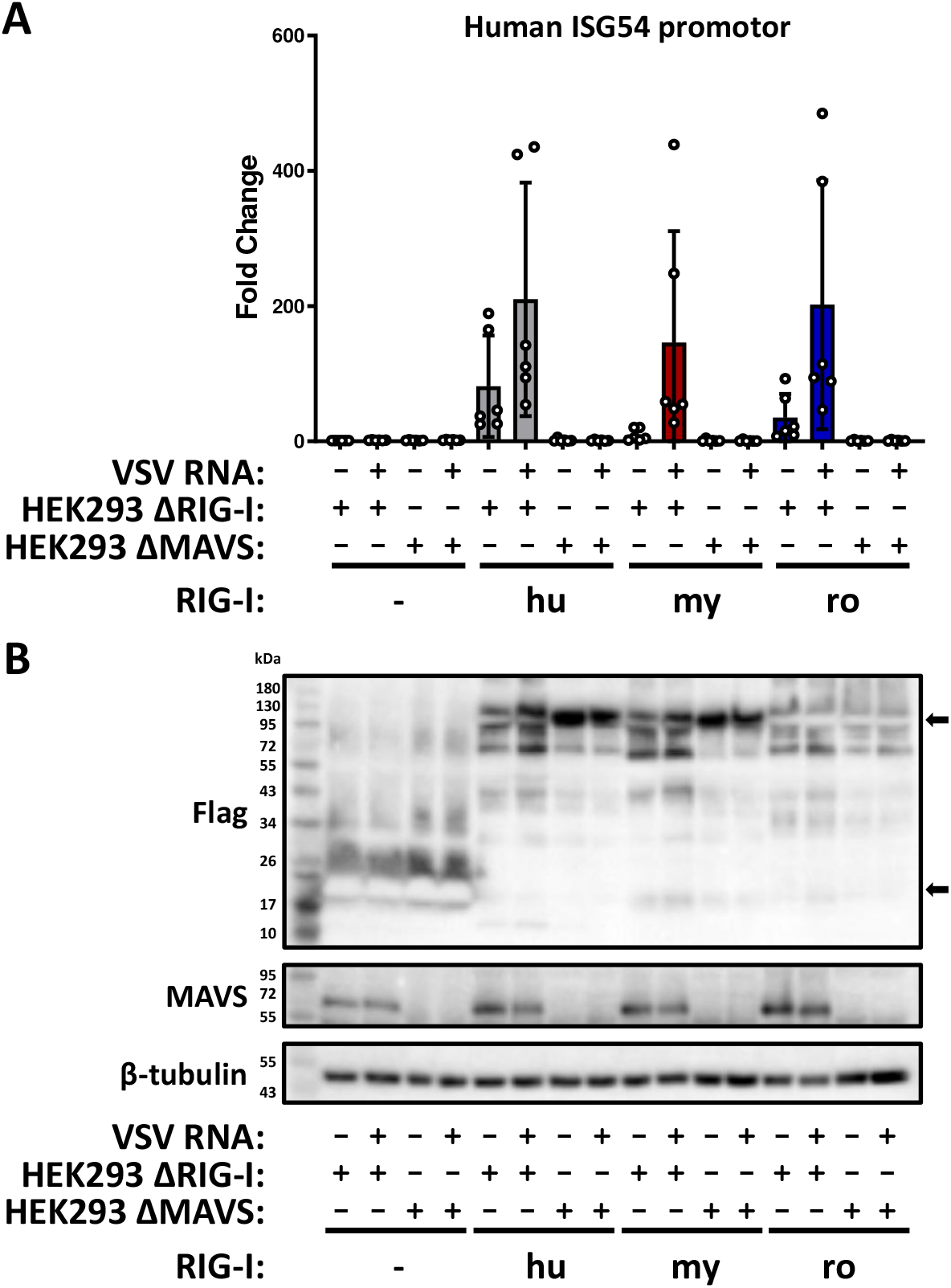
MAVS-dependent antiviral signalling by human and bat RIG-I orthologs. **A)** HEK293 ΔRIG-I or ΔMAVS cells were transfected, stimulated and assayed as described for figure 3. The graph shows the results from six independent replicates. **B)** Immunoblot analysis was performed with antibodies against the indicated antigens. The expected size of the 3xFlag-ΔMx control (-) and RIG-I proteins are indicated with arrows. Representative data from five independent experiments are shown.

### Bat RIG-I signaling via MAVS

Upon recognition of 5’ppp-dsRNA, RIG-I undergoes a conformational change to expose the CARDs for interaction with MAVS which, in turn, initiates the antiviral signalling. To investigate whether the bat RIG-I orthologs are also signaling via MAVS, we compared their ability to activate the ISG54-promotor reporter in HEK293 ΔRIG-I and HEK293 ΔMAVS cells. The cells with the respective genotype were transfected with the expression constructs and stimulated with VSV RNA as described. RIG-I-dependent ISG54-promotor activation was only detected in the presence of MAVS (Fig. 5A), with comparable expression levels of the Flag-tagged RIG-I orthologs (Fig. 5B). Thus, the bat RIG-I, like the human ortholog, are signaling via MAVS.

### *RNA ligand* interaction

We investigated the interaction of the bat RIG-I orthologs with RNA. Firstly, we performed pulldowns with the biotin-labelled dsRNA analog polyI:C (HMW) that is bound to streptavidin-beads. Cell lysates from HEK293 ΔRIG-I cells expressing either Flag-tagged control or RIG-I proteins (Fig. 6A, left panel) were mixed with the polyI:C-coated beads. To make sure that the binding was specific, we also incubated the lysates with either empty beads or added free VSV RNA as a competitor. After incubation, washing and elution, the eluates were subjected to SDS-PAGE and immunoblot detection of the Flag epitope tag (Fig. 6A, right panel). None of the RIG-I orthologs bound unspecifically to the empty beads, but all exhibited a binding to the poly I:C-coated beads which could be efficiently outcompeted by free VSV RNA.

**Figure 6.**
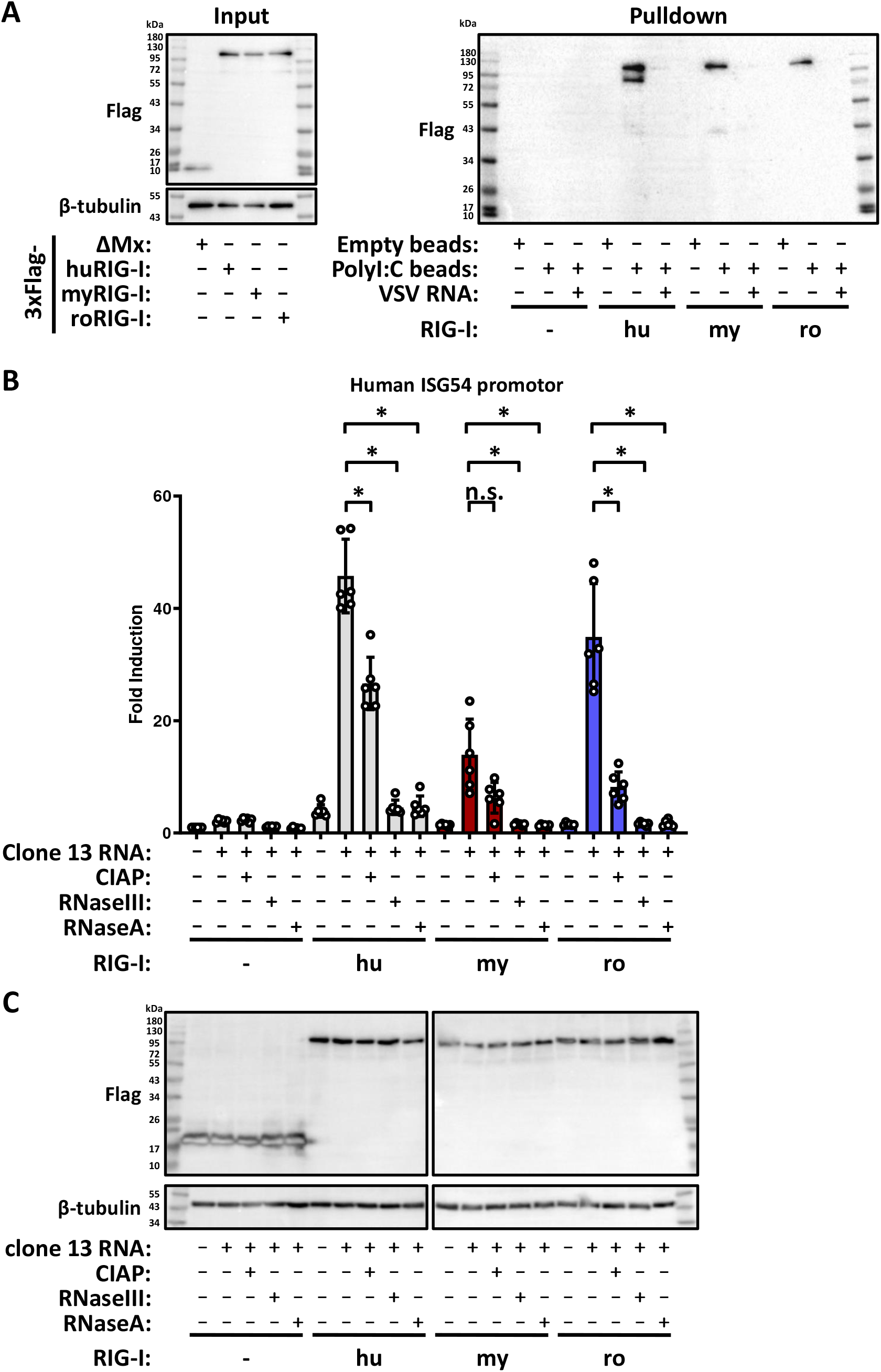
Involvement of double stranded RNA and 5’-phophorylation in RIG-I activation. **A)** PolyI:C pulldown. Cell lysates from HEK293 ΔRIG-I cells transfected with plasmids encoding 3xFlag-ΔMx, hu-, my- or roRIG-I were incubated with magnetic beads coupled, or not, with HMW-polyI:C. Possible binding of the respective RIG-I to the polyI:C were competed with free VSV genomic RNA. After incubation and elution, immunoblot analysis was performed for the Flag epitope tag. Representative data from three independent experiments are shown. **B)** HEK293 ΔRIG-I cells were transfected with cDNA constructs for 3xFlag-ΔMx, hu-, my- or roRIG-I and stimulated with VSV genomic RNA that was pretreated with the different enzymes as indicated. After 24 h the cells were harvested and the firefly/*Renilla* luciferase activity measured. The graph shows the results from six independent replicates. **C)** Immunoblot analysis using antibodies against the indicated antigens. Representative data from six independent experiments are shown. n.s.: non-significant, * p<0,05.

Next, we investigated whether the activation of the RIG-I orthologs depends on the 5’-triphosphate moiety and the double-strandedness of a viral RNA. To this aim, genomic RNA isolated from Rift Valley fever virus clone 13 particles were either mock, Calf Intestinal Alkaline Phosphatase (CIAP, 5’-phosphatase), RNaseIII (double-stranded RNA (dsRNA) specific endoribonuclease) or RNaseA treated. The RNAs were then transfected into HEK293 ΔRIG-I cells that had been transfected with the RIG-I expressing plasmids and the ISG54-reporter plasmid. After 24 h of incubation, ISG54 promotor activity was measured. Pre-treatment of the RNA with CIAP more than halved the induction by all RIG-Is, compared to mock treated RNA, and pretreatment with either of the two RNases reduced the induction level to background, irrespective of the particular RIG-I (Fig. 6B to C). Together with the polyI:C pulldown, the above results indicate that bat RIG-I, like human and mouse RIG-I (27, 28), are specifically binding dsRNA and that full activation by viral genome RNA requires both a 5’-triphosphate and double stranded RNA.

### IFN induction by SARS-CoV-2 can be enabled by RIG-I orthologs

*Rhinolophus* “megabats” (suborder *Yinpterochiroptera*) are proposed to be reservoirs of SARS-coronaviruses including the pandemic SARS-CoV-2 (3). Infection of human lung epithelial cells with SARS-CoV-2 induces a certain level of type I IFN (53), which was mostly found to be mediated by MDA5, a PRR that is structurally and functionally related to RIG-I (54-56). However, also RIG-I was found to be involved in IFN and cytokine induction by SARS-CoV-2, either directly via antiviral signalling (57, 58), or indirectly by controlling viral RNA synthesis down to non-inducing levels (59). We tested the involvement of our bat RIG-I orthologs for their ability to sense SARS-CoV-2 infection. Transcomplementation experiments in human ACE2-HEK293 ΔRIG-I cells demonstrated that the RIG-I orthologs of humans and of *R. aegyptiacus* are indeed capable of inducing IFN-beta mRNA synthesis in response to SARS-CoV-2 (Fig. 7 A, top left panel). RIG-I of *M. daubentonii*, by contrast, was not reacting to SARS-CoV-2. *R. aegyptiacus* RIG-I, but not RIG-I from the two other species, also raised mRNA levels of the chemokine CXCL10, but just by a factor below 2 (Fig. 7 A, top right panel). For both human and *R. aegyptiacus* RIG-I orthologs, the cytokine mRNA induction levels by SARS-CoV-2 were approximately 5 to more than 10 fold lower than for RVFV clone 13. This is expected, as unlike SARS-CoV-2 the positive control virus clone 13 is devoid of any IFN antagonistic factor. No differences in SARS or RVFV RNA (Fig. 7 A, bottom panels) or protein levels (Fig. 7 B) were observed between the negative control (3xFlag-ΔMx) and the RIG-I expressing samples (Fig. 7 B), and expression of all three RIG-I orthologs was confirmed (Fig. 7 C and suppl. Fig. S5). Thus, the RIG-I orthologs of both humans and the “megabat” *R. aegyptiacus*, but not of “microbat” *M. daubentonii*, enable innate immune sensing of SARS-CoV-2. However, expression of the various RIG-I orthologs also induced MDA5 (Fig 7 C). Although the levels of MDA5 induction were comparatively low, overexpressing RIG-I orthologs could not entirely clarify whether the IFN response to SARS-CoV-2 was owed to the respective RIG-I orthologs, or rather to the endogenous MDA5 they are inducing.

**Figure 7.**
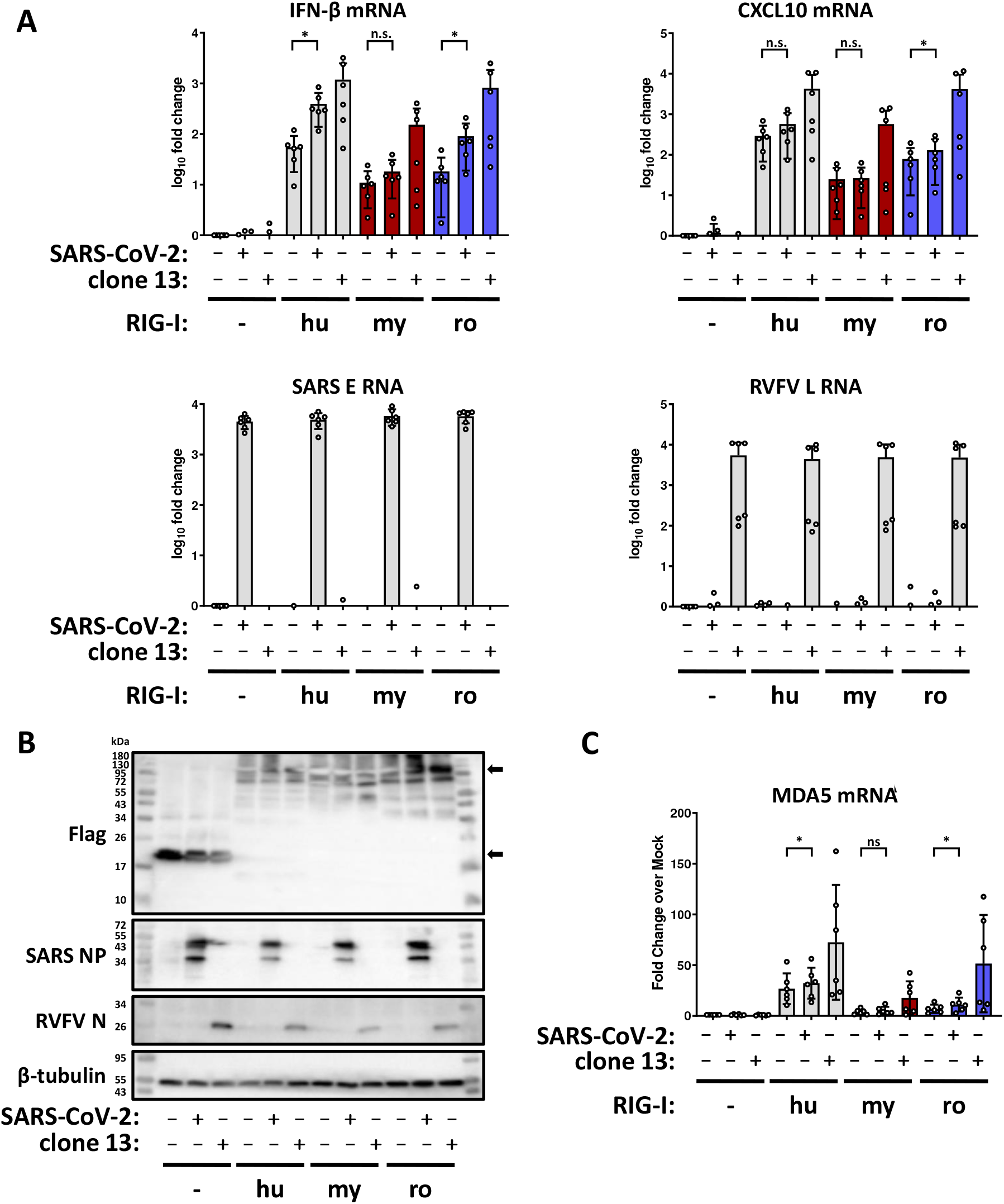
Innate immune response to SARS-CoV-2 in cells overexpressing RIG-I orthologs. **A)** ACE2-HEK293 ΔRIG-I cells were transfected with the indicated plasmids for 24 h, followed by infection with either SARS-CoV-2 or RVFV clone 13 (MOI 1). After 16 h, total RNA was isolated and used for detection of IFN-β and CXCL10 mRNA, SARS E RNA and RVFV L RNA by RT-qPCR. The graph shows the results from six independent replicates. **B)** Immunoblot analysis with antibodies against the indicated antigens. Representative data from six independent experiments are shown. **C)** RT-qPCR analysis for MDA5 mRNA with the samples used in **A)**. n.s.: non-significant, * p<0,05

### MDA5 is dispensable for SARS-CoV-2 innate immune sensing

To investigate whether the RIG-I-dependent IFN induction by SARS-CoV-2 could be indirectly mediated, we abrogated MDA5 expression by siRNA treatment before transfection of ACE2-HEK293 ΔRIG-I cells with the two SARS-CoV-2-reactive RIG-I orthologs (human and *R. aegyptiacus*). As shown in figure 8A, the IFN induction in response to SARS-CoV-2 occurred irrespective of whether cells were treated with the control siRNA or with the MDA5 siRNA, as long as human or *R. aegyptiacus* RIG-I were present. The previously seen (low) effect of SARS-CoV-2 on CXCL10 induction could not be reproduced in the siRNA-transfected cells (Fig. S6A). Again, even in the RIG-I expressing cells the MDA5 levels were comparatively low and suppressed by the specific siRNA as expected (Fig. 8B and S6B to D). Moreover, viral RNA levels including those of SARS-CoV-2 were not influenced by the overexpressed RIG-I orthologs (Fig. S6 E and F). Thus, in our system MDA5 seems not to contribute to the RIG-I-mediated IFN induction by SARS-CoV-2.

**Figure 8.**
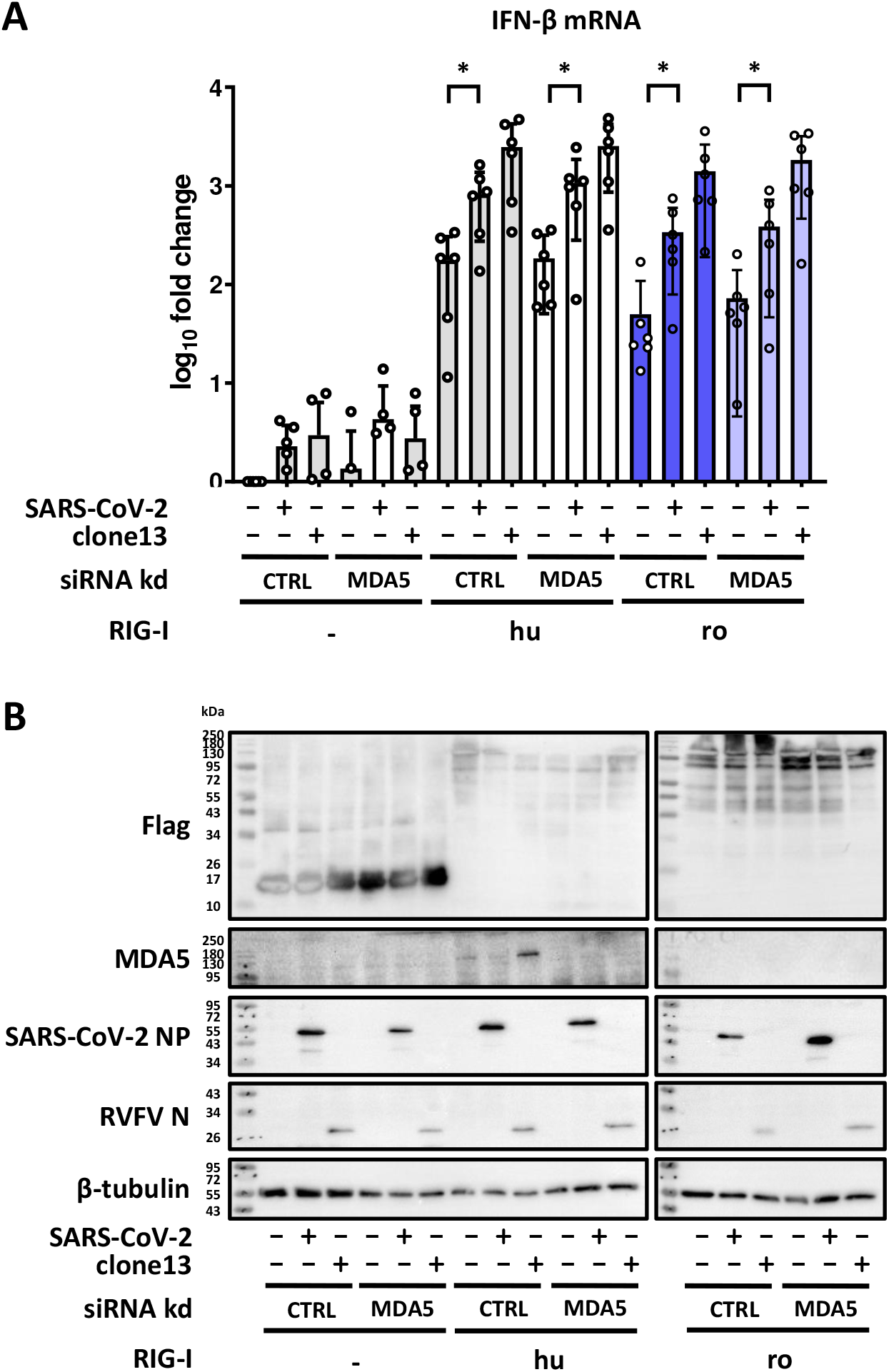
Innate immune response to SARS-CoV-2 is RIG-I dependent. **A)** Expression of endogenous MDA5 was siRNA-knocked down in the ACE2-HEK293 ΔRIG-I cells. The cells were then transfected with the indicated plasmids for 24 h, followed by infection with either SARS-CoV-2 or RVFV clone 13 (MOI 1). After 16 h, total RNA was isolated and used for detection of IFN-β mRNA by RT-qPCR. The graph shows the results from six independent replicates. **B)** Immunoblot analysis with antibodies against the indicated antigens. Representative data from six independent experiments are shown. * p<0,05.

Overall, these experiments demonstrate that the RIG-I orthologs of *R. aegyptiacus* and humans, but not of *M. daubentonii*, are indeed capable of sensing infection by SARS-CoV-2.

## DISCUSSION

RIG-I is responsible for one of the earliest steps of the innate immune response, the recognition of intruding viral RNA and the induction of antiviral IFNs (28, 34). Since bats are a major reservoir for zoonotic viruses and suspected to have a special IFN system (1-3), we set out to deep-characterize RIG-I orthologs from representatives of both the “microbats” (*M. daubentonii*, suborder *Yangochiroptera*) and the “megabats” (*R. aegyptiacus*, suborder *Yinpterochiroptera*), and compare them to human RIG-I.

In terms of gene expression, we could stimulate mRNA synthesis of the bat RIG-I orthologs in parental cells by type I IFN treatment as well as by virus infection, results that are in line with previous findings (24, 35-40, 42). The amino acid sequences derived from cloned RIG-I cDNAs were more than 90% similar between each other and compared to human RIG-I (with the notable exception of small indels) and the predicted domain structure was obviously conserved.

Our functional characterization of the *M. daubentonii* and *R. aegyptiacus* RIG-I orthologs indicate that bat RIG-Is are not substantially different from the human counterpart. All three RIG-I orthologs were capable of binding dsRNA, are in their function partially dependent on the 5’ triphosphate RNA end, are signaling via MAVS, and can trigger the induction of IFN and other cytokines in response to viral RNA or virus infection. In the overexpression experiments, neither protein levels nor dsRNA binding or 5’triphosphate end dependency were substantially different. Nonetheless, the RIG-I of *M. daubentonii*, exhibited in most settings a somewhat reduced activity, whereas *R. aegyptiacus* RIG-I tended more often to perform comparably to human RIG-I. Importantly, the reduced activity by *M. daubentonii* RIG-I occurred in both a human and a mouse cell background. When parental bat cells were stimulated with a RIG-I-inducing virus, by contrast, neither the antiviral response in total nor IFN induction *per se* were much different between the *M. daubentonii* and *R. aegyptiacus* RIG-I. Thus, *Myotis* RIG-I may not possess a systematically lower antiviral signalling capability, but rather need a cofactor that is only present or only fitting in parental cells and regulates a signalling step downstream of RNA recognition. Further studies are however necessary to clarify this.

Unlike humans, bats have to endure extreme changes in body temperature when they switch between hibernation, sleep, or flight (51, 60). Nonetheless we did not observe any species-specific differences regarding temperature-dependent activity of the respective RIG-Is. The two bat and the human RIG-I orthologs showed RNA-responsive activity at 37°C and at 39 °C, while at 30°C there was only background activity (which was highest for the human ortholog). Although we cannot exclude a dominant influence of the human cell background, the observations are in agreement with a publication on the reduced and delayed IFN system activity at lower temperatures in human, primate and mouse systems (52), and with the report that hibernating bats elicit very little immune responses to a fungal pathogen (60).

The involvement of RIG-I in IFN induction by SARS-CoV-2 is still not entirely clarified. While most reports concluded that MDA5 rather than RIG-I is the relevant PRR (54-56), two studies reported it is additionally RIG-I-dependent (57, 58). In our hands, both human and *R. aegyptiacus* RIG-I enabled HEK293 ΔRIG-I cells to induce IFN in response to SARS-CoV-2 infection, and in an MDA5-independent manner. RIG-I seems therefore to be another PRR sensing SARS-CoV-2 RNA under certain conditions. Besides this, it is interesting that the RIG-I of *R. aegyptiacus*, but not of *M. daubentonii* is reacting to SARS-CoV-2, as *Rousettus* belongs to the same suborder *Yinpterochiroptera* as *Rhinolophus*, the major host of SARS-coronaviruses (3).

The antiviral IFN system of bats has attracted great attention as it might be the key to understand the special role of these animals as virus reservoirs. The results presented are corroborating previous studies and predictions on the high similarity of bat RIG-I orthologs with those of other mammals regarding sequence, domain structure and regulation. Our functional data on dsRNA binding and the dependency on 5’ triphosphate ends, MAVS and temperature, as well as on the recognition of SARS-CoV-2 infection do not indicate substantial differences between the RIG-Is of humans and bats. This insight complements recent observations that the antiviral IFN effectors Tetherin and PKR of bats are basically functioning the same way as their human counterparts (23, 25). On the other hand, however, compared to humans some bats encode multiple copies of the genes for Tetherin and PKR as well as for IFNs (20, 23-25). Thus, it could be speculated that the IFN system of bats has functionally conserved key factors, but is probably enforced by increased copy numbers of antiviral genes.

In any case, it is hoped that the data and tools we generated will enable further insights into the role of the antiviral IFN system for the unique capacity of bats to harbour a huge range of different human pathogenic viruses without falling prey to them.

## MATERIALS & METHODS

### Cells and viruses

A549, MyDauNi (MyDauNi/2c), Vero E6, BHK, HEK293 ΔRIG-I or ΔMAVS, ACE2-HEK293 ΔRIG-I, MEF RIG-I-/-, were maintained in DMEM supplemented with 17,8 mg/L L-alanine, 0,7 g/L glycine, 75 mg/L L-glutamic acid, 25 mg/L L-proline, 0,1 mg/L biotin, 25 mg/L hypoxanthine, and 3,7 g/L sodium bicarbonate, 10% fetal calf serum (FCS), 2 mM glutamine, 120 U/ml penicillin, and 100 g/ml streptomycin while Ro6E-J were cultivated in DMEM (Gibco, 21969035) supplemented with 10% fetal calf serum (FCS), 2 mM glutamine, 120 U/ml penicillin, and 100 g/ml streptomycin (Gibco, 10378016) and non-essential amino acids (Gibco, 11140050). Stocks of RVFV clone 13, LACV wt, LACV ΔNSs and VSV were grown on BHK cells while SARS-CoV-2 (München-1.2/2020/984 (B.1) (61)) was propagated on VeroE6 cells, infected with an MOI of 0,001, for 72 h (RVFV/LACV/SARS-CoV-2) or 48 h for VSV. The respective stock were then titrated on Vero E6 for SARS-CoV-2, using an overlay medium containing MEM (Gibco, 21935028) supplemented with 10% fetal bovine serum, 2 mM glutamine, 120 U/ml penicillin, and 100 g/ml streptomycin (Gibco, 10378016) and 1,5 % Avicell (62). RVFV clone 13, LACV wt/ΔNSs and SARS-CoV-2 was incubated for 72 h while VSV was incubated for 24 h. The overly was then removed and the cells fixed and stained using staining solution (0.75% crystal violet, 3.75% formaldehyde, 20% ethanol, 1% methanol).

### RT-qPCR analyses

The cells were infected as indicated in the figure. After the indicted time point, total cellular RNA was isolated using RNeasy Mini kit (Qiagen, Cat No./ID: 74106) according to manufacturer’s instructions. A total of 100 ng isolated RNA was used for cDNA synthesis using PrimeScript High Fidelity RT-PCR Kit (Takara, R022B). RT-qPCR was performed using TB Green® Premix Ex Taq™ (Tli RNase H Plus) (Takara, RR420B) according to manufacturer’s instructions on an Applied biosystems StepOnePlus machine. For detection of human transcripts, QuantiTect Primer Assay (Qiagen) against 18S ribosomal RNA (QT00199367), IFN-β (QT00203763), RIG-I (QT00040509), CXCL10 (QT01003065), MxA (QT00090895), OAS1 (QT00099134) and MDA5 (QT00033789) were used. For the detection of full-length transcript sequences for *M. daubentonii*, we used an available *de novo* transcriptome assembly based on bulk RNA-Seq data (SRR8062281-SRR8062299) from our previous study (24). In short, we applied an ensemble approach as described in (63) combining the output of different transcriptome assembly tools and used the final assembly (available at https://osf.io/x9kad) for detecting full-length transcripts for *M. daubentonii*. For *R. aegyptiacus*, we used publicly available genome and annotation data to obtain transcript sequences (GCF_014176215.1). Primers were designed using Primer3Plus (https://primer3plus.com/cgi-bin/dev/primer3plus.cgi) and ordered from Eurofins. For detection of *M. daubentonii* 18S ribosomal RNA: fwd 5’ AAACGGCTACCACATCCAAG 3’ and rev 5’ CCTCCAATGGATCCTCGTTA 3’, IFN-β: fwd 5’ AAAGCAGCAATTCAGCCTGT 3’ and rev 5’ CTGCTGGAGCATCTCGTACA 3’, RIG-I: fwd 5’ GGAAAACCACAACCTGCACT 3’ and rev 5’ ACTCTTTGGTCTGGGGTGTG 3’, CXCL10: fwd 5’ TTTTCTGCCTCATCCTTCTGA 3’ and rev 5’ TGGACAAGATGGACTTGCAG 3’, IFN-λ3: fwd 5’ CACATCCACTCCAAGCTTCA 3’ and rev 5’ TCAGCGACACATCTCAGGTC 3’, Mx1: fwd 5’ CAGAGGGAGAGGGCTTTCTT 3’ and rev 5’ TCTGCTGGTTCTCCTTTATTTG 3’ and OAS1: fwd 5’ AGCCATTGACACCATCTGCA 3’ and rev 5’ CTCTTGCTGACATGCTTCCA 3’ while for *R. aegyptiacus* 18S ribosomal RNA: fwd 5’ CGCGGTTCTATTTTGTTGGT 3’ and rev 5’ AGTCGGCATCGTTTATGGTC 3’, IFN-β: fwd 5’ ATTGCCTCAAGGACAGGATG 3’ and rev 5’ TTCAGTTTCTCCAGGGCTGT 3’, RIG-I: fwd 5’ CAAAAGCACAAGTGAAGCCT 3’ and rev 5’ TTGTCGGTAGTCCGTGATTC 3’, CXCL10: fwd 5’ TCAACCTGTTAATCCAAAGTCC 3’ and rev 5’ CCTTTCCTTGCTAATTGCTTTC 3’, IFN-λ3: fwd 5’ ACCTCCACCACTGGCTGT 3’ and rev 5’ AATGGCAACACGTTTCAGGT 3’, Mx1: fwd 5’ TCGGCTGTTTACCAAAATCC 3’ and rev 5’ CCAGGGTTTTGATTTGCTGT 3’ and OAS1: fwd 5’ CTATGCTTGGGAACGTGGAT 3’ and rev 5’ GGCCAACTCTGTGAGTCTCC 3’ were used. RVFV L segment RNA and SARS E RNA were detected with PrimeDirect™ Probe RT-qPCR Mix (Takara, RR650A) according to manufacturer’s instructions using RVFV L primers fwd 5’ TGAAAATTCCTGAGACACATGG 3’, rev 5’ ACTTCCTTGCATCATCTGATG 3’ and probe 5’ 6FAM-CAATGTAAGGGGCCTGTGTGGACTTGTG-BHQ1 3’ (64) or the SARS-CoV-2 E primers fwd 5’ ACAGGTACGTTAATAGTTAATAGCGT 3’, rev 5’ ATATTGCAGCAGTACGCACACA 3’ and probe 5’ FAM-ACACTAGCCATCCTTACTGCGCTTCG-BBQ 3’ (65). The results are presented as the ΔΔCT-value using 18 ribosomal RNA as internal control (66).

### Virus titration

Supernatants from cells that had been infected with LACV wt or LACV ΔNS were collected and cleared by centrifugation at 800×g for 5 min. The supernatants were titrated on Vero E6 with an overlay medium containing MEM (Gibco, 21935028) supplemented with 10% fetal bovine serum, 2 mM glutamine, 120 U/ml penicillin, and 100 g/ml streptomycin (Gibco, 10378016) and 1,5 % Avicel (FMC BioPolymer, (62)). After 48 h of incubation, the medium was removed and the cells washed with PBS before fixing with PBS-4% paraformaldehyde (Roth, 0335-4) for 24 h at 4° C. Then, the fixed cells were again washed with PBS and permeabilized with PBS-0,1% Triton X-100 (Sigma, T9284-500ML) for 20 min. The cells were again washed and then incubated for 16 h at 4°C with Anti-LACV N (1:1000, kind gift from Georg Kochs, University or Freiburg, Germany) in TBS-T buffer containing 1 % non-fat milk powder. After washing the cells with PBS, they were incubated with secondary antibody IRDye800 conjugated anti-Rabbit (1:10000, Rockland, 611-132-122) and DRAQ5 (1:10000, eBioscience, 65-0880-92) in TBS-T-1 % milk powder at room temperature. Finally, the wells were washed first with PBS and then with H_2_O, and the fluorescence signals were detected and the foci counted using the Odyssey instrument (LI-COR).

### cDNA Cloning

Primers for cloning human, *M. daubentonii* and *R. aegyptiacus* RIG-I were generated using the In-Fusion cloning tool (https://www.takarabio.com/learning-centers/cloning/primer-design-and-other-tools) and ordered from Eurofins Genomics. The full-length sequences of RIG-I (DDX58) from *M. daubentonii* was assembled from previous data (24) while the human and *R. aegyptiacus* RIG-I sequences were available as GenBank entries (NM_014314.4 and XM_016130339.2, respectively). Primers were designed for cloning the cDNAs into the vector pI.18, with a 5’ 3xFlag-tag sequence added to the forward primer. Human RIG-I fwd: 5’ TGA CAC GAT CGG ATC CAT GGA CTA CAA AGA CCA TGA CGG TGA TTA TAA AGA TCA TGA TAT CGA TTA CAA GGA TGA CGA TGA CAA GAC CAC CGA GCA GCG ACG C 3’ and rev 5’ TCT AGA ATT CCT CGA GTC ATT TGG ACA TTT CTG CTG GA 3’; *M. daubentonii* and *R. aegyptiacus* RIG-I: fwd 5’ TGA CAC GAT CGG ATC CAT GGA CTA CAA AGA CCA TGA CGG TGA TTA TAA AGA TCA TGA TAT CGA TTA CAA GGA TGA CGA TGA CAA GAC GGC CGA GGA GCG GCG G 3’, *M. daubentonii* rev 5’ TCT AGA ATT CCT CGA GTC ATT TGG ACA TTT CTG CTG GAT C 3’, *R. aegyptiacus* rev 5’ TCT AGA ATT CCT CGA GTC ATT TGG GCA TTT CTG CAA CAT CG 3’. cDNAs were generated from RNAs isolated from A549, MyDauNi and Ro6E-J cells that were treated for 16 h with 1000 U/ml IFN-α (B/D) (PBL Assay Science). The RNAs were isolated using RNeasy Mini kit (Qiagen, Cat No./ID: 74106), and 1 µg was used for cDNA synthesis with the PrimeScript High Fidelity RT-PCR Kit (Takara, R022B). Of the cDNA, 2 µl were used as PCR template for amplification with the KOD Polymerase (Calbiochem, 71086-3). The PCR products were cloned into the pI.18 vector, digested with *Bam*HI (NEB, R3136S) and *Kpn*I (NEB, R3142S), using the In-Fusion kit (Takara, 638911). The ligated product was transformed into Stellar competent cells accompanying the In-Fusion kit, and spread onto agar plates at 37° C for 16h before colonies were picked for DNA isolation. Correctness of the inserts was confirmed by sequencing.

### Bioinformatics analysis

The full-length sequences of the cloned RIG-Is for *Homo sapiens, M. daubentonii*, and *R. aegyptiacus* were aligned to the respective published sequence for human (NM_014314.4) and *R. aegyptiacus* (XM_016130339.2) RIG-I, respectively, while the *M. daubentonii* sequence was compared to our assembled sequence from our previous study (24) (transcriptome available at https://osf.io/x9kad). The respective bat RIG-Is were completely conserved while the human RIG-I had a silent mutation at position 2709 (A to G) and 2760 (A to T). The DNA sequences were then *in silico* translated using the Expasy translation tool (https://web.expasy.org/translate/) and the resulting amino acid sequences aligned using T-Coffee (http://tcoffee.crg.cat/apps/tcoffee/do:regular) (67) and visualized with Boxshade (https://embnet.vital-it.ch/software/BOX_form.html). The respective domains were manually assigned based on Kolakofsky *et al*. (68) and confirmed by running the respective amino acid sequence in the SMART database (http://smart.embl-heidelberg.de/).

### Generation of ACE2-HEK293 ΔRIG-I cells

For efficient infection studies with SARS-CoV-2, we transduced HEK293 ΔRIG-I cells with a lentivirus expressing human ACE2 (hACE2, EC:3.4.17.23) under the control of an EF-1α promoter. For this purpose, we used the lentiviral expression system ViraPower (Thermofisher, K4975-00) and the transgene packaging vector pEGIP-Puro (pEGIP was a gift from Linzhao Cheng, Addgene Plasmid 26777; http://n2t.net/addgene:26777), which couples the expression of the transgene to the expression of the selection marker using an intra ribosomal entry site (IRES) (DOI: 10.1016/j.stem.2009.05.023). The pEGIP-derived packaging vector pEGIP-hACE2 was generated by homologous recombination using the NEBuilder kit (NEB, E5520). The vector backbone was amplified using oligonucleotides with appropriate hACE2 sequence overhangs, namely pEGIP_hACE2_fwd 5’ TGATGATGTTCAGACCTCCTTTAGCCGCCCCCCCCCTCTC 3’ and pEGIP_hACE2_rev 5’ GCCAGGAAGAGCTTGACATCGATATCAAGCTTACCTAGC 3’. The hACE2 gene was amplified using the oligonucleotides ACE2_fwd 5’ ATGTCAAGCTCTTCCTGGCTC 3’ and ACE2_rev 5’CTAAAAGGAGGTCTGAACATCATC 3’ from a plasmid containing the hACE2 mRNA. VSV-G pseudotyped lentiviral vector particles were produced in 293T cells by transfection of 1 µg pEGIP-hACE2 (transgene packaging vector), 0.8 µg pLP1 (gag, pol, and rev expression), 0.6 µg pLP2 (rev expression), and 0.3 µg pLP3 (VSV-G expression). Three days after transfection, the supernatant was harvested, sterile filtered through a 0.2 µm syringe adapter, and used to transduce 6 × 106 HEK293 ΔRIG-I cells. Two days after transduction, the supernatant was removed, cells were harvested by trypsinization and re-seeded in 10-fold dilution series in DMEM containing 1 µg/ml puromycin. Single grown cell foci were scraped out of the cell culture dish after one week, cells were then separated by trypsinization and further selected by limited 10-fold dilution, yielding single cellular clones. After verification of hACE2 expression by immunofluorescence assays, the clonal cell line HEK293 ACE2 ΔRIG-I was expanded from one clone, cryo-preserved, and used for the experiments.

### RIG-I transcomplementation assays

A total of 5×10^4^ HEK293 ΔRIG-I, HEK293 ΔMAVS, ACE2-HEK293 ΔRIG-I, or MEF RIG-I^-/-^ cells were seeded in 24-well plates. After 24 h the cells were transfected with 0,25 µg µg of either pI.18-3xFlag-ΔMx (negative control), pI.18-3xFlag-huRIG-I, pI.18-3xFlag-myRIG-I or pI.18-3xFlag-roRIG-I together with transfection control pLR-SV40-*Renilla* (Promega) and either of the reporter plasmids ISG54-Luc (69), p125-Luc (70) or pGL4.10 Rousettus IFN-ßp (71), as indicated in the respective figure, using the GeneJammer (Agilent, 204130) transfection reagent. Half of the plasmid transfected wells were stimulated by either transfecting 250 ng/well genomic VSV RNA or RVFV clone 13 RNA, using the Endofectin (GeneCopoeia, EF013) transfection reagent, or infecting the cells with RVFV clone 13 (MOI 10), as indicated in the respective figure. After 16 h of incubation the cells were lysed in Passive Lysis Buffer (Promega) and the firefly and *Renilla* luciferase activity measured using the Dual Luciferase assay kit (Promega, E1960) and a TriStar^2^ Multimode Reader LB942 (Berthold technologies). The data are presented as fold over unstimulated pI.18-3xFlag-ΔMx (CTRL) with firefly reporter values normalised to *Renilla* reporter control.

### Assay for RNA dependence of RIG-I

A total of 125 ng genomic RVFV clone 13 RNA was either mock treated or incubated with 40 U CIAP (Promega, M1821), 2 U RNaseIII (Ambion, AM2290) or 2 µg RNaseA (Ambion, AM2269) for 16 h at 37 °C. An aliquot of 25 ng/well of the treated RNAs were the mixed with either pI.18-3xFlag-ΔMx (CTRL), pI.18-3xFlag-huRIG-I, pI.18-3xFlag-myRIG-I or pI.18-3xFlag-roRIG-I (25 ng/well each) together with transfection control pLR-SV40-*Renilla* (Promega) (50 ng/well) and the reporter plasmids ISG54-Luc (250 ng/well) (69) and transfected into HEK293 ΔRIG-I cells that were grown overnight in 24-well plates as described for the transcomplementation assay, using Endofectin (GeneCopoeia, EF013). After an additional 24 h of incubation, firefly and *Renilla* luciferase activities were measured and processed as described for the transcomplementation assay.

### Isolation of genomic *VSV and RVFV clone 13 RNA*

Genomic RNAs were isolated from VSV or RVFV clone 13 particles by PEG800-precipitation and phenol-chloroform extraction as described (72). Briefly, the supernatant from a T175 flask infected with VSV or RVFV clone 13 (MOI 0,001) was collected and cleared by centrifugation at 800×g for 5 min, after 48 h (VSV) or 72 h (RVFV) of incubation. The cleared supernatant was then mixed with 10,8 ml PEG8000 buffer (30% (w/v) PEG 8000, 10 mM Tris-HCl (pH 7,5), 1 mM EDTA, 100 mM NaCl) and 1,5 mL 5M NaCl followed by incubated for 1 h at 4°C with head-over-tail rotation. The solution was then centrifugated at 4600×g for 1 h at 4°C and the resulting pellet dissolved in 1,5 ml peqGOLD TriFast (peqlab, 30-2030). After vortexing and incubation at room temperature for 5 min, a phase-separation was performed by centrifugation at 4600×g for 11 min. The upper aqueous phase was collected and mixed with 700 µl chloroform (Roth, 7331) and glycogen (Roche, 10 901 393 001). The RNA was then precipitated at -20°C for 16 h and pelleted by centrifugation at 4600×g for 33 min at 4°C. The pellet was washed twice with 70% Ethanol (Roth, 9065.4) and then dissolved in H_2_O to a final concentration of 250 ng/µl.

### Immunoblot analysis

Cell lysates were mixed 4:1 with 4× sample buffer (143 mM Tris-HCl (pH 6,8), 28,6% Glycerol, 5,7 % SDS, 4.3 mM Bromphenol Blue and 5% 2-mercaptoethanol) supplemented with phosphatase (Calbiochem, 524625) and protease (Roche, 04 693 116 001) inhibitors. The lysates were boiled for 10 min, separated by SDS-PAGE, and then blotted onto polyvinylidene difluoride membranes. The membranes were blocked with 5% (wt/vol) non-fat dry milk powder in TBS-T and probed with primary antibodies against the following targets: Flag (Sigma, F3165, 1:2000, mouse, monoclonal), anti-β-tubulin (Abcam, ab6046, 1:1000, rabbit, polyclonal), MAVS (Alexis, ALX-210-929, 1:1000, rabbit, polyclonal), RVFV N (1:2000, rabbit, polyclonal (serum), kind gift from Alejandro Brun), MDA5 (CellSignaling, 5321S, 1:1000, rabbit, monoclonal) and SARS-CoV-2 NP (biomol, 200-401-A50, 1:2000, rabbit, polyclonal). The secondary antibodies were peroxidase-conjugated anti-mouse IgG (Thermo Fisher, catalog no. 31430, 1:40,000, goat, polyclonal) and peroxidase-conjugated anti-rabbit IgG (Thermo Fisher, catalog no. 31460, 1:40,000, goat, polyclonal). Western blot signals were detected on a Chemidoc (Bio-Rad) using SuperSignal West Femto maximum sensitivity substrate (Thermo Scientific, catalog no. 34096).

### Poly I:C pull down

A total of 2×10^5^ HEK293 ΔRIG-I cells per well were seeded in 6-well plates. After 24 h the cells were transfected with 100 ng/well of either pI.18-3xFlag-ΔMx (CTRL), pI.18-3xFlag-huRIG-I, pI.18-3xFlag-myRIG-I or pI.18-3xFlag-roRIG-I using GeneJammer (Agilent, 204130) transfection reagent. After 16 h of incubation the cells were scraped off in PBS and pelleted by centrifugation at 800×g for 5 min. The cells were then lysed in lysis buffer (0,5% Triton X-100 (Sigma, T9284-500ML), 1X Protease inhibitor (c0mplete, Roche, 4693116001) in PBS) for 10 min at 4°C. Cell debris were removed by centrifugation at 15000×g, 10 min, 4°C and the supernatants used for pull down. As input from each reaction, 5% of the volume was put aside. Dynabeads M-270 Streptavidin (Invitrogen, 65305) (250 µg/IP) was coupled to 100 ng/IP Poly(I:C) (HMW) Biotin (Invivogen) according to manufacturer’s instructions. After washing, the beads were re-suspended in the cell lysates and incubated at 4°C with head over tail rotation for 16 h. To compete for RIG-I binding, 10 µg of VSV RNA was added to the lysate before incubation. After incubation the beads were washed three times in lysis buffer and the bound fraction eluted in 0,5% Triton X-100 in PBS mixed 4:1 in 4× sample buffer supplemented with phosphatase and protease inhibitors. The samples were then boiled and the supernatant used for immunoblot analysis.

### SARS-CoV-2 infection

HEK293 ΔRIG-I-ACE2 cells, 5×10^4^ cells/well, were seeded in 24-well plates treated with Poly-D-lysine hydrobromide (Sigma, P6407-5MG) and reverse transfected with 250 ng/well of either pI.18-3xFlag-ΔMx, pI.18-3xFlag-huRIG-I, pI.18-3xFlag-myRIG-I or pI.18-3xFlag-roRIG-I using GeneJammer transfection reagent (Agilent technologies, 204131). After 24 h of incubation, the cells were infected with SARS-CoV-2 or RVFV clone 13 at an MOI of 1 under BSL3 conditions. After 16h of incubation the medium was removed, the cells washed with PBS and the samples taken for RT-qPCR or immunoblotting.

### siRNA treatment

ACE2-HEK293 ΔRIG-I cells were seeded at 5×10E4 cells per well in 24-well plates that had been treated with Poly-D-lysine hydrobromide (Sigma, P6407-5MG) and reverse transfected with 50 nM/well of control (CTRL) or MDA5 siRNA (Qiagen, FlexiTube Genesolution Cat. No.: 1027280 and 1027416) using Lipofectamine RNAiMAX (Thermo Fisher, 13778075). After 4 h of incubation the medium was exchanged and the cells transfected with 250 ng/well of either pI.18-3xFlag-ΔMx, pI.18-3xFlag-huRIG-I, pI.18-3xFlag-myRIG-I or pI.18-3xFlag-roRIG-I using GeneJammer transfection reagent (Agilent technologies, 204131). After 24 h of incubation, the cells were infected with SARS-CoV-2 or RVFV clone 13 at an MOI of 1 as described above. After 16h of incubation the medium was removed, the cells washed with PBS and the samples taken for RT-qPCR or immunoblotting.

### Statistical analysis

Two-tailed paired t-tests were done using Graphpad Prism, based on the logarithmic values for viral growth assays while all other comparisons were made on fold-induction over control values.

## FIGURE LEGENDS

**Suppl. Figure S1. Continuation of Fig. 1A to show type I Interferon competence of micro- and megabat cells lines**. cDNAs from cells described in figure 1 were analysed for mRNA expression of CXCL10, MxA/Mx1 and OAS1 by RT-qPCR of the respective species. The graphs show the fold induction over mock data points, with mean values and standard deviations from three independent replicates.

**Suppl. Figure S2. Nucleotide sequence alignment for human, micro- and megabat RIG-I**.

Symbols are as described for figure 2.

**Suppl. Figure S3. SMART result for human, micro- and megabat RIG-I**. The amino acid sequences for our cloned human and bat RIG-I orthologs were analyzed using the SMART database. Pictures of the output are shown.

**Suppl. Figure S4. Transcomplementation of RIG-I-deficient mouse cells by bat RIG-I orthologs. A)** MEF RIG-I^-/-^ cells were transfected with plasmids encoding CTRL, hu-, my- or roRIG-I together with firefly-luciferase expressing plasmids under the control of the *Rousettus* IFN-β promotor and *Renilla*-luciferase under SV40-control promotor. After 24 h the cells were stimulated by infection with RVFV clone 13 (MOI 10) and 16 h later analysed for firefly/*Renilla* luciferase activities. The graphs show the fold induction over CTRL data points, with mean values and standard deviations from three independent replicates. **B)** Immunoblot analysis with antibodies against the indicated antigens. Representative data from three independent experiments are shown. * p<0,05.

**Suppl. Figure S5. Extended control panel for figure 7. A)** cDNAs from cells described in figure 7A were analysed by RT-qPCR for mRNA expression of RIG-I of the respective species. The graphs show the fold induction over mock data points, with mean values and standard deviations from three independent replicates. **B)** Quantification of the immunoblot analyses for viral nucleoproteins, normalized to ß-tubulin, as shown in figure 7B.

**Suppl. Figure S6. Extended control panel for figure 8. A)** to **F)** cDNAs from cells described in figure 8 were analysed by RT-qPCR for the presence of the indicated RNA sequences. n.s.: non-significant, * p<0.05.

## ACKNOWLEDGMENTS

We thank Besim Berisha, Anna Hoffbauer and Patrick Schmerer for technical assistance, and Alejandro Brun and Georg Kochs kindly providing antisera. Work in the F.W. laboratory is funded by the Deutsche Forschungsgemeinschaft (DFG) SFB 1021 (project number 197785619) and SPP 1596 (grant We 2616/7-2), and by the RAPID consortium of the Bundesministerium für Bildung und Forschung (BMBF; 01KI1723E and 01KI20158). The work of C.D. was supported by grants received from the BMBF (01KI2006A), and the DFG (DR 772/10-2). M.A.M. received funds from the Volkswagen Foundation (AZ93345).

## REFERENCES

1. Brook CE, Dobson AP. 2015. Bats as ‘special’ reservoirs for emerging zoonotic pathogens. Trends Microbiol 23:172–80.

2. Luis AD, Hayman DTS, O’Shea TJ, Cryan PM, Gilbert AT, Pulliam JRC, Mills JN, Timonin ME, Willis CKR, Cunningham AA, Fooks AR, Rupprecht CE, Wood JLN, Webb CT. 2013. A comparison of bats and rodents as reservoirs of zoonotic viruses: are bats special? Proceedings of the Royal Society B-Biological Sciences 280.

3. Van Brussel K, Holmes EC. 2021. Zoonotic disease and virome diversity in bats. Curr Opin Virol 52:192–202.

4. Hu B, Zeng LP, Yang XL, Ge XY, Zhang W, Li B, Xie JZ, Shen XR, Zhang YZ, Wang N, Luo DS, Zheng XS, Wang MN, Daszak P, Wang LF, Cui J, Shi ZL. 2017. Discovery of a rich gene pool of bat SARS-related coronaviruses provides new insights into the origin of SARS coronavirus. Plos Pathogens 13.

5. Zhou H, Ji J, Chen X, Bi Y, Li J, Wang Q, Hu T, Song H, Zhao R, Chen Y, Cui M, Zhang Y, Hughes AC, Holmes EC, Shi W. 2021. Identification of novel bat coronaviruses sheds light on the evolutionary origins of SARS-CoV-2 and related viruses. Cell doi:10.1016/j.cell.2021.06.008.

6. Leroy EM, Kumulungui B, Pourrut X, Rouquet P, Hassanin A, Yaba P, Delicat A, Paweska JT, Gonzalez JP, Swanepoel R. 2005. Fruit bats as reservoirs of Ebola virus. Nature 438:575–576.

7. Towner JS, Pourrut X, Albarino CG, Nkogue CN, Bird BH, Grard G, Ksiazek TG, Gonzalez JP, Nichol ST, Leroy EM. 2007. Marburg virus infection detected in a common African bat. PLoS One 2:e764.

8. Teeling EC, Vernes SC, Davalos LM, Ray DA, Gilbert MTP, Myers E, Bat KC. 2018. Bat Biology, Genomes, and the Bat1K Project: To Generate Chromosome-Level Genomes for All Living Bat Species. Annu Rev Anim Biosci 6:23–46.

9. Irving AT, Ahn M, Goh G, Anderson DE, Wang LF. 2021. Lessons from the host defences of bats, a unique viral reservoir. Nature 589:363–370.

10. Ahn M, Cui J, Irving AT, Wang LF. 2016. Unique Loss of the PYHIN Gene Family in Bats Amongst Mammals: Implications for Inflammasome Sensing. Sci Rep 6:21722.

11. Ahn M, Anderson DE, Zhang Q, Tan CW, Lim BL, Luko K, Wen M, Chia WN, Mani S, Wang LC, Ng JHJ, Sobota RM, Dutertre CA, Ginhoux F, Shi ZL, Irving AT, Wang LF. 2019. Dampened NLRP3-mediated inflammation in bats and implications for a special viral reservoir host. Nat Microbiol 4:789–799.

12. Banerjee A, Rapin N, Bollinger T, Misra V. 2017. Lack of inflammatory gene expression in bats: a unique role for a transcription repressor. Sci Rep 7:2232.

13. Prescott J, Guito JC, Spengler JR, Arnold CE, Schuh AJ, Amman BR, Sealy TK, Guerrero LW, Palacios GF, Sanchez-Lockhart M, Albarino CG, Towner JS. 2019. Rousette Bat Dendritic Cells Overcome Marburg Virus-Mediated Antiviral Responses by Upregulation of Interferon-Related Genes While Downregulating Proinflammatory Disease Mediators. mSphere 4.

14. Xie J, Li Y, Shen X, Goh G, Zhu Y, Cui J, Wang LF, Shi ZL, Zhou P. 2018. Dampened STING-Dependent Interferon Activation in Bats. Cell Host Microbe 23:297–301 e4.

15. Banerjee A, Zhang X, Yip A, Schulz KS, Irving AT, Bowdish D, Golding B, Wang LF, Mossman K. 2020. Positive Selection of a Serine Residue in Bat IRF3 Confers Enhanced Antiviral Protection. iScience 23:100958.

16. Clayton E, Munir M. 2020. Fundamental Characteristics of Bat Interferon Systems. Front Cell Infect Microbiol 10:527921.

17. Fuchs J, Holzer M, Schilling M, Patzina C, Schoen A, Hoenen T, Zimmer G, Marz M, Weber F, Muller MA, Kochs G. 2017. Evolution and Antiviral Specificities of Interferon-Induced Mx Proteins of Bats against Ebola, Influenza, and Other RNA Viruses. J Virol 91.

18. Hawkins JA, Kaczmarek ME, Muller MA, Drosten C, Press WH, Sawyer SL. 2019. A metaanalysis of bat phylogenetics and positive selection based on genomes and transcriptomes from 18 species. Proc Natl Acad Sci U S A 116:11351–11360.

19. Zhang G, Cowled C, Shi Z, Huang Z, Bishop-Lilly KA, Fang X, Wynne JW, Xiong Z, Baker ML, Zhao W, Tachedjian M, Zhu Y, Zhou P, Jiang X, Ng J, Yang L, Wu L, Xiao J, Feng Y, Chen Y, Sun X, Zhang Y, Marsh GA, Crameri G, Broder CC, Frey KG, Wang LF, Wang J. 2013. Comparative analysis of bat genomes provides insight into the evolution of flight and immunity. Science 339:456–60.

20. Pavlovich SS, Lovett SP, Koroleva G, Guito JC, Arnold CE, Nagle ER, Kulcsar K, Lee A, Thibaud-Nissen F, Hume AJ, Muhlberger E, Uebelhoer LS, Towner JS, Rabadan R, Sanchez-Lockhart M, Kepler TB, Palacios G. 2018. The Egyptian Rousette Genome Reveals Unexpected Features of Bat Antiviral Immunity. Cell 173:1098–1110 e18.

21. Zhou P, Cowled C, Mansell A, Monaghan P, Green D, Wu L, Shi Z, Wang LF, Baker ML. 2014. IRF7 in the Australian black flying fox, Pteropus alecto: evidence for a unique expression pattern and functional conservation. PLoS One 9:e103875.

22. Zhou P, Tachedjian M, Wynne JW, Boyd V, Cui J, Smith I, Cowled C, Ng JH, Mok L, Michalski WP, Mendenhall IH, Tachedjian G, Wang LF, Baker ML. 2016. Contraction of the type I IFN locus and unusual constitutive expression of IFN-alpha in bats. Proc Natl Acad Sci U S A 113:2696–701.

23. Hayward JA, Tachedjian M, Johnson A, Irving AT, Gordon TB, Cui J, Nicolas A, Smith I, Boyd V, Marsh GA, Baker ML, Wang LF, Tachedjian G. 2022. Unique Evolution of Antiviral Tetherin in Bats. J Virol 96:e0115222.

24. Holzer M, Schoen A, Wulle J, Muller MA, Drosten C, Marz M, Weber F. 2019. Virus- and Interferon Alpha-Induced Transcriptomes of Cells from the Microbat Myotis daubentonii. iScience 19:647–661.

25. Jacquet S, Culbertson M, Zhang C, El Filali A, De La Myre Mory C, Pons JB, Filippi-Codaccioni O, Lauterbur ME, Ngoubangoye B, Duhayer J, Verez C, Park C, Dahoui C, Carey CM, Brennan G, Enard D, Cimarelli A, Rothenburg S, Elde NC, Pontier D, Etienne L. 2022. Adaptive duplication and genetic diversification of protein kinase R contribute to the specificity of bat-virus interactions. Sci Adv 8:eadd7540.

26. Lazear HM, Schoggins JW, Diamond MS. 2019. Shared and Distinct Functions of Type I and Type III Interferons. Immunity 50:907–923.

27. Liu G, Gack MU. 2020. Distinct and Orchestrated Functions of RNA Sensors in Innate Immunity. Immunity 53:26–42.

28. Yoneyama M, Onomoto K, Jogi M, Akaboshi T, Fujita T. 2015. Viral RNA detection by RIG-I-like receptors. Curr Opin Immunol 32:48–53.

29. Kowalinski E, Lunardi T, McCarthy AA, Louber J, Brunel J, Grigorov B, Gerlier D, Cusack S. 2011. Structural basis for the activation of innate immune pattern-recognition receptor RIG-I by viral RNA. Cell 147:423–35.

30. Luo D, Ding SC, Vela A, Kohlway A, Lindenbach BD, Pyle AM. 2011. Structural insights into RNA recognition by RIG-I. Cell 147:409–22.

31. Rehwinkel J, Tan CP, Goubau D, Schulz O, Pichlmair A, Bier K, Robb N, Vreede F, Barclay W, Fodor E, Reis e Sousa C. 2010. RIG-I detects viral genomic RNA during negative-strand RNA virus infection. Cell 140:397–408.

32. Weber M, Gawanbacht A, Habjan M, Rang A, Borner C, Schmidt AM, Veitinger S, Jacob R, Devignot S, Kochs G, Garcia-Sastre A, Weber F. 2013. Incoming RNA virus nucleocapsids containing a 5’-triphosphorylated genome activate RIG-I and antiviral signaling. Cell Host Microbe 13:336–46.

33. Weber M, Sediri H, Felgenhauer U, Binzen I, Banfer S, Jacob R, Brunotte L, Garcia-Sastre A, Schmid-Burgk JL, Schmidt T, Hornung V, Kochs G, Schwemmle M, Klenk HD, Weber F. 2015. Influenza virus adaptation PB2-627K modulates nucleocapsid inhibition by the pathogen sensor RIG-I. Cell Host Microbe 17:309–319.

34. Weber M, Weber F. 2014. RIG-I-like receptors and negative-strand RNA viruses: RLRly bird catches some worms. Cytokine Growth Factor Rev 25:621–8.

35. Cowled C, Baker ML, Zhou P, Tachedjian M, Wang LF. 2012. Molecular characterisation of RIG-I-like helicases in the black flying fox, Pteropus alecto. Dev Comp Immunol 36:657–64.

36. He X, Korytar T, Zhu Y, Pikula J, Bandouchova H, Zukal J, Kollner B. 2014. Establishment of Myotis myotis cell lines--model for investigation of host-pathogen interaction in a natural host for emerging viruses. PLoS One 9:e109795.

37. Li J, Zhang G, Cheng D, Ren H, Qian M, Du B. 2015. Molecular characterization of RIG-I, STAT-1 and IFN-beta in the horseshoe bat. Gene 561:115–23.

38. Holzer M, Krahling V, Amman F, Barth E, Bernhart SH, Carmelo VAO, Collatz M, Doose G, Eggenhofer F, Ewald J, Fallmann J, Feldhahn LM, Fricke M, Gebauer J, Gruber AJ, Hufsky F, Indrischek H, Kanton S, Linde J, Mostajo N, Ochsenreiter R, Riege K, Rivarola-Duarte L, Sahyoun AH, Saunders SJ, Seemann SE, Tanzer A, Vogel B, Wehner S, Wolfinger MT, Backofen R, Gorodkin J, Grosse I, Hofacker I, Hoffmann S, Kaleta C, Stadler PF, Becker S, Marz M. 2017. Differential transcriptional responses to Ebola and Marburg virus infection in bat and human cells (vol 6, 34589, 2016). Scientific Reports 7.

39. Sarkis S, Lise MC, Darcissac E, Dabo S, Falk M, Chaulet L, Neuveut C, Meurs EF, Lavergne A, Lacoste V. 2018. Development of molecular and cellular tools to decipher the type I IFN pathway of the common vampire bat. Dev Comp Immunol 81:1–7.

40. Tarigan R, Shimoda H, Doysabas KCC, Ken M, Iida A, Hondo E. 2020. Role of pattern recognition receptors and interferon-beta in protecting bat cell lines from encephalomyocarditis virus and Japanese encephalitis virus infection. Biochem Biophys Res Commun 527:1–7.

41. Tarigan R, Katta T, Takemae H, Shimoda H, Maeda K, Iida A, Hondo E. 2021. Distinct interferon response in bat and other mammalian cell lines infected with Pteropine orthoreovirus. Virus Genes 57:510–520.

42. Glennon NB, Jabado O, Lo MK, Shaw ML. 2015. Transcriptome Profiling of the Virus-Induced Innate Immune Response in Pteropus vampyrus and Its Attenuation by Nipah Virus Interferon Antagonist Functions. J Virol 89:7550–66.

43. Mostajo NF, Lataretu M, Krautwurst S, Mock F, Desiro D, Lamkiewicz K, Collatz M, Schoen A, Weber F, Marz M, Holzer M. 2020. A comprehensive annotation and differential expression analysis of short and long non-coding RNAs in 16 bat genomes. NAR Genom Bioinform 2:lqz006.

44. Schoen A, Lau S, Verbruggen P, Weber F. 2020. Elongin C Contributes to RNA Polymerase II Degradation by the Interferon Antagonist NSs of La Crosse Orthobunyavirus. J Virol 94.

45. Blakqori G, Delhaye S, Habjan M, Blair CD, Sanchez-Vargas I, Olson KE, Attarzadeh-Yazdi G, Fragkoudis R, Kohl A, Kalinke U, Weiss S, Michiels T, Staeheli P, Weber F. 2007. La Crosse bunyavirus nonstructural protein NSs serves to suppress the type I interferon system of mammalian hosts. J Virol 81:4991–9.

46. Hess RD, Weber F, Watson K, Schmitt S. 2012. Regulatory, biosafety and safety challenges for novel cells as substrates for human vaccines. Vaccine 30:2715–2727.

47. Schmid S, Mordstein M, Kochs G, Garcia-Sastre A, Tenoever BR. 2010. Transcription factor redundancy ensures induction of the antiviral state. J Biol Chem 285:42013–22.

48. Kainulainen M, Habjan M, Hubel P, Busch L, Lau S, Colinge J, Superti-Furga G, Pichlmair A, Weber F. 2014. Virulence factor NSs of rift valley fever virus recruits the F-box protein FBXO3 to degrade subunit p62 of general transcription factor TFIIH. J Virol 88:3464–73.

49. Garcia-Lopez MA, Sancho D, Sanchez-Madrid F, Marazuela M. 2001. Thyrocytes from autoimmune thyroid disorders produce the chemokines IP-10 and Mig and attract CXCR3+ lymphocytes. J Clin Endocrinol Metab 86:5008–16.

50. Haller O, Staeheli P, Schwemmle M, Kochs G. 2015. Mx GTPases: dynamin-like antiviral machines of innate immunity. Trends Microbiol 23:154–63.

51. Burbank RC, Young JZ. 1934. Temperature changes and winter sleep of bats. J Physiol 82:459–67.

52. Lane WC, Dunn MD, Gardner CL, Lam LKM, Watson AM, Hartman AL, Ryman KD, Klimstra WB. 2018. The Efficacy of the Interferon Alpha/Beta Response versus Arboviruses Is Temperature Dependent. mBio 9.

53. Banerjee A, El-Sayes N, Budylowski P, Jacob RA, Richard D, Maan H, Aguiar JA, Demian WL, Baid K, D’Agostino MR, Ang JC, Murdza T, Tremblay BJ, Afkhami S, Karimzadeh M, Irving AT, Yip L, Ostrowski M, Hirota JA, Kozak R, Capellini TD, Miller MS, Wang B, Mubareka S, McGeer AJ, McArthur AG, Doxey AC, Mossman K. 2021. Experimental and natural evidence of SARS-CoV-2-infection-induced activation of type I interferon responses. iScience 24:102477.

54. Rebendenne A, Valadao ALC, Tauziet M, Maarifi G, Bonaventure B, McKellar J, Planes R, Nisole S, Arnaud-Arnould M, Moncorge O, Goujon C. 2021. SARS-CoV-2 triggers an MDA-5-dependent interferon response which is unable to control replication in lung epithelial cells. J Virol doi:10.1128/JVI.02415-20.

55. Sampaio NG, Chauveau L, Hertzog J, Bridgeman A, Fowler G, Moonen JP, Dupont M, Russell RA, Noerenberg M, Rehwinkel J. 2021. The RNA sensor MDA5 detects SARS-CoV-2 infection. Sci Rep 11:13638.

56. Yin X, Riva L, Pu Y, Martin-Sancho L, Kanamune J, Yamamoto Y, Sakai K, Gotoh S, Miorin L, De Jesus PD, Yang CC, Herbert KM, Yoh S, Hultquist JF, Garcia-Sastre A, Chanda SK. 2021. MDA5 Governs the Innate Immune Response to SARS-CoV-2 in Lung Epithelial Cells. Cell Reports 34.

57. Arai Y, Yamanaka I, Okamoto T, Isobe A, Nakai N, Kamimura N, Suzuki T, Daidoji T, Ono T, Nakaya T, Matsumoto K, Okuzaki D, Watanabe Y. 2023. Stimulation of interferon-beta responses by aberrant SARS-CoV-2 small viral RNAs acting as retinoic acid-inducible gene-I agonists. iScience 26:105742.

58. Thorne LG, Reuschl AK, Zuliani-Alvarez L, Whelan MVX, Turner J, Noursadeghi M, Jolly C, Towers GJ. 2021. SARS-CoV-2 sensing by RIG-I and MDA5 links epithelial infection to macrophage inflammation. Embo Journal 40.

59. Yamada T, Sato S, Sotoyama Y, Orba Y, Sawa H, Yamauchi H, Sasaki M, Takaoka A. 2021. RIG-I triggers a signaling-abortive anti-SARS-CoV-2 defense in human lung cells. Nat Immunol 22:820–828.

60. Field KA, Sewall BJ, Prokkola JM, Turner GG, Gagnon MF, Lilley TM, Paul White J, Johnson JS, Hauer CL, Reeder DM. 2018. Effect of torpor on host transcriptomic responses to a fungal pathogen in hibernating bats. Mol Ecol doi:10.1111/mec.14827.

61. Rothe C, Schunk M, Sothmann P, Bretzel G, Froeschl G, Wallrauch C, Zimmer T, Thiel V, Janke C, Guggemos W, Seilmaier M, Drosten C, Vollmar P, Zwirglmaier K, Zange S, Wolfel R, Hoelscher M. 2020. Transmission of 2019-nCoV Infection from an Asymptomatic Contact in Germany. N Engl J Med 382:970–971.

62. Matrosovich M, Matrosovich T, Garten W, Klenk HD. 2006. New low-viscosity overlay medium for viral plaque assays. Virol J 3:63.

63. Holzer M, Marz M. 2019. De novo transcriptome assembly: A comprehensive cross-species comparison of short-read RNA-Seq assemblers. Gigascience 8.

64. Bird BH, Bawiec DA, Ksiazek TG, Shoemaker TR, Nichol ST. 2007. Highly sensitive and broadly reactive quantitative reverse transcription-PCR assay for high-throughput detection of Rift Valley fever virus. J Clin Microbiol 45:3506–13.

65. Corman VM, Landt O, Kaiser M, Molenkamp R, Meijer A, Chu DK, Bleicker T, Brunink S, Schneider J, Schmidt ML, Mulders DG, Haagmans BL, van der Veer B, van den Brink S, Wijsman L, Goderski G, Romette JL, Ellis J, Zambon M, Peiris M, Goossens H, Reusken C, Koopmans MP, Drosten C. 2020. Detection of 2019 novel coronavirus (2019-nCoV) by real-time RT-PCR. Euro Surveill 25.

66. Livak KJ, Schmittgen TD. 2001. Analysis of relative gene expression data using real-time quantitative PCR and the 2(T)(-Delta Delta C) method. Methods 25:402–408.

67. Notredame C, Higgins DG, Heringa J. 2000. T-Coffee: A novel method for fast and accurate multiple sequence alignment. J Mol Biol 302:205–17.

68. Kolakofsky D, Kowalinski E, Cusack S. 2012. A structure-based model of RIG-I activation. RNA 18:2118–27.

69. Paulson M, Press C, Smith E, Tanese N, Levy DE. 2002. IFN-Stimulated transcription through a TBP-free acetyltransferase complex escapes viral shutoff. Nature Cell Biology 4:140–147.

70. Yoneyama M, Suhara W, Fukuhara Y, Fukuda M, Nishida E, Fujita T. 1998. Direct triggering of the type I interferon system by virus infection: activation of a transcription factor complex containing IRF-3 and CBP/p300. EMBO J 17:1087–95.

71. Papies J, Sieberg A, Ritz D, Niemeyer D, Drosten C, Muller MA. 2022. Reduced IFN-ss inhibitory activity of Lagos bat virus phosphoproteins in human compared to Eidolon helvum bat cells. PLoS One 17:e0264450.

72. Habjan M, Andersson I, Klingstrom J, Schumann M, Martin A, Zimmermann P, Wagner V, Pichlmair A, Schneider U, Muhlberger E, Mirazimi A, Weber F. 2008. Processing of Genome 5′ Termini as a Strategy of Negative-Strand RNA Viruses to Avoid RIG-I-Dependent Interferon Induction. Plos One 3.

